# The hidden intricacies of aquaporins: Remarkable details in a common structural scaffold

**DOI:** 10.1101/2022.03.28.486021

**Authors:** Nikolaus Gössweiner-Mohr, Christine Siligan, Kristyna Pluhackova, Linnea Umlandt, Sabina Köfler, Natasha Trajkovska, Andreas Horner

**Affiliations:** Institute of Biophysics, Johannes Kepler University Linz, Gruberstr. 40, 4020 Linz, Austria; Stuttgart Center for Simulation Science, Cluster of Excellence EXC 2075, Universitätstr. 32, University of Stuttgart, 70569 Stuttgart, Germany

## Abstract

Evolution turned aquaporins (AQPs) into the most efficient facilitators of passive water flow through cell membranes at no expense of solute discrimination. In spite of a plethora of solved AQP structures, many structural details remain hidden. Here, by combining extensive sequence- and structural-based analysis of a unique set of 20 non-redundant high-resolution structures and molecular dynamics simulations of 4 representatives, we identify key aspects of AQP stability, gating, selectivity, pore geometry and oligomerization, with a potential impact on channel functionality. We challenge the general view of AQPs possessing a continuous open water pore and depict that AQPs selectivity is not exclusively shaped by pore lining residues but also by the relative arrangement of transmembrane helices. Moreover, our analysis reveals that hydrophobic interactions constitute the main determinant of protein thermal stability. Finally, we establish a novel numbering scheme of the conserved AQP scaffold facilitating direct comparison and prediction of potential structural effects of e.g. disease-causing mutations. Additionally, our results pave the way for the design of optimized AQP water channels to be utilized in biotechnological applications.

## 1. Introduction

Aquaporins (AQPs), part of a larger family of major intrinsic proteins, are one of the best studied protein families with currently 20 non-redundant high-resolution structures (≤3,70 Å) solved. Since their first discovery in 1992 by Peter Agre and coworkers (1, 2), thirteen different types of aquaporins (AQP0-12) were discovered in mammals (3). The narrow AQP pores combine enormous permeability, conducting water in a single-file manner close to the diffusion limit of water in bulk, with exceptional selectivity (4). A subset of AQPs, the aquaglyceroporins (AQGPs), paralogs of AQPs, are also able to conduct glycerol and other small neutral solutes (5, 6). Bacteria also express members of AQPs and AQGPs, generally functioning with one copy of each paralog and, interestingly, some lacking both. Unicellular eukaryotes and fungi follow a similar pattern, with a clear division between AQPs or AQGPs and a heterogeneous distribution in the number of copies of each paralog in the different genera (7). So far, no archaea has been found that possesses both paralogs concurrently. Plants exhibit the highest AQP diversity, with five main subfamilies (plasma membrane intrinsic proteins (PIPs), tonoplast intrinsic proteins (TIPs), nodulin-26 like intrinsic proteins (NIPs), small basic intrinsic proteins (SIPs) and X intrinsic proteins (XIPs)), which are each further divided into subgroups (7). Furthermore, in primitive plant species, two additional subfamilies, GLPF-like intrinsic proteins (GIPs) and hybrid intrinsic proteins (HIPs), have been found (8). However, the full diversity of AQ(G)Ps is still not represented by the numerous high-resolution structures, as exemplified by only three plant aquaporin structures and none of the unorthodox AQ(G)Ps (represented by AQP11 and 12 in mammals).

Their critical involvement in cellular water homeostasis and great selectivity renders AQ(G)Ps important in several key areas: (i) AQ(G)Ps are expressed in a wide variety of tissues throughout the mammalian body, where they play a role in an extensive range of physiological functions (6), including water/salt homeostasis, exocrine fluid secretion and epidermal hydration. Due to their important tasks throughout the body, AQ(G)Ps are involved in various human diseases, including glaucoma, cancer, epilepsy, and obesity (9, 10). Mutations in their primary sequence cause genetic diseases like nephrogenic diabetes insipidus, congenital cataracts and keratoderma (11). Therefore, AQ(G)Ps represent potential drug targets (10, 12, 13). Moreover, (ii) AQ(G)Ps fulfill pivotal functions in plants, where they also participate in the regulation of cellular water homeostasis (14), thus steering transpiration sensitivity to soil drying as well as to high atmospheric vapor pressure deficit (15). Hence, they represent the perfect target to address abiotic stresses, like drought, through genetic engineering (16). Drought is one of the major threats of agricultural production worldwide and therefore, research efforts have focused on the development of drought resistant crops in the past years. To reach this goal, it is important to explore the underlying molecular mechanisms as well as the inter-related pathways and signaling networks through which AQ(G)Ps induce drought tolerance (16). Additionally, (iii) due to their structural stability and easy handling, AQPs are candidates for building blocks of next generation filter membranes (17–28) and are already used as a benchmark for newly designed artificial water channels in terms of permeability and selectivity (29). Thus, concepts of solute and solvent flux through narrow membrane channels can be directly transferred to this challenging and emerging field of material science (30–37) and used to optimize their performance.

AQ(G)Ps exhibit two constriction sites, the aromatic/arginine selectivity filter (ar/R), with its aromatic residue(s) and the positively charged arginine, and the dual asparagine-proline-alanine (NPA) motive, with the structurally opposing half-helices creating a positive dipole moment, both of which are involved in proton rejection (38–40). Progress in our understanding of mechanisms underlaying AQ(G)P functionality was mainly achieved by structural investigations (41–58), sequence comparisons (8), non-quantitative *in vitro* assays (38), yeast complementation assays (38, 59) and *in silico* studies (39, 60, 61). It was generally believed that the amino acid composition of the ar/R-region determines the differences in the specificity of AQPs and AQGPs, primarily by affecting the pore size (38, 60, 61). However, several studies pointed out that solute specificity cannot be explained by differences in pore size alone (60, 62, 63). One of the latest studies emphasized that substrate discrimination in water channels depends on a complex interplay between solute, pore size as well as polarity, and that channel pore size determines exclusion properties but not solute selectivity (64). Besides water, selected AQ(G)Ps are shown to facilitate a wide variety of neutral molecules across lipid bilayers (65), like hydrogen peroxide (59, 66), metalloids (67, 68), monocarboxylic acid species (69), ammonia (58) and lactate/lactic acid (70, 71). In individual cases, the substrate selectivity was linked to a specific residue, e.g. His63 in helix 2 of *At*Tip2;1 is crucial for ammonia specificity (58), or substitution of Ile254 with Met at the periplasmic end of helix 6 in *Hv*PIP2;3 impairs transport of CO_2_, while having no impact on water permeability (72). Alternatively, the presence of Cys residues lining the water pore was suggested to be connected to the ability to transport H_2_O_2_ (73). However, the general molecular determinants enabling the permeability of single substrates across their highly conserved tertiary and quaternary structure remain largely elusive. This may partly be due to the lack of quantitative methods to estimate neutral solute permeabilities.

Quantitative unitary water permeabilities *p_f_* of AQPs, summarized in Figure 1, are rare and span almost one order of magnitude. Currently, diverse methods are used to obtain *p_f_*, delivering sometimes more and sometimes less coherent results (74–78). Only recently, driven by methodological advancements (4, 76, 79, 80), the major determinants of single-file transport of water through membrane channels were found to be the number of H-bonds water molecules form with pore lining residues (4), positive charges at the pore mouth (81), channel gating (77) and potentially the shape of the vestibule (82). Thereby, high *p_f_* is intricately linked to a low activation energy of water transport (83).

**Figure 1.**
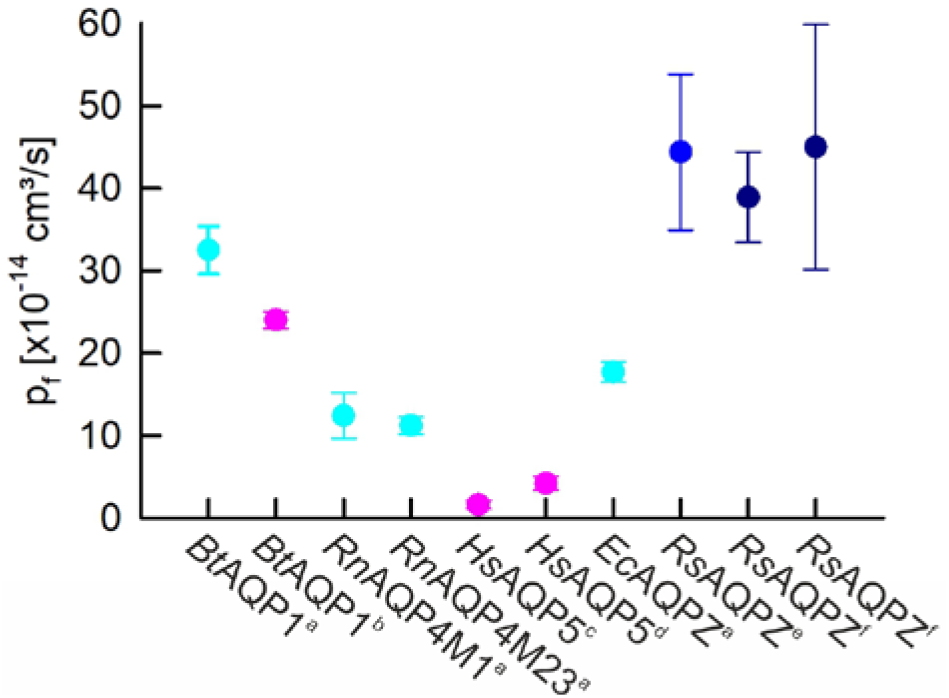
Published quantitative single channel water permeabilities p_f_ of tetrameric AQPs at diverse temperatures. p_f_ values are included if they are based on accurate channel counting and P_f_ estimation. ^a^Stopped-flow light scattering and fluorescence correlation spectroscopy (FCS) counting in Escherichia coli polar lipid extract (PLE) at 5°C (4, 81), ^b^Micropipette aspiration technique and FCS at room temperature (RT) (74), ^c^Polarized cell monolayer with FCS at RT (84), ^d^Single cell swelling using laser scanning microscopy (LSM) and FCS at RT (84), ^e^Stopped-flow light scattering and FCS counting in a 4:1 phosphatidylcholine/ phosphatidylserine (PC/PS) mixture at 10°C, ^f^Stopped-flow light scattering and FCS counting in PMOXA-PDMS-PMOXA at 15°C. Values are temperature-color coded (5°C – cyan, 10°C – blue, 15°C – dark blue, RT – pink).

Although many excellent papers and reviews on AQ(G)Ps have been published, covering their evolution (7, 85, 86), potential roles in disease (9–13), AQ(G)P gating (3, 87–90), biotechnological applications (28, 91–93) and their protein-protein interactions (94, 95), to name a few, a thorough analysis elucidating their subtle structural differences and determining the molecular basis for solute selectivity, protein gating, and protein stability, is missing so far. Even recent structural papers and reviews analyze and compare only small sub-sets of AQ(G)P structures, which does not educe patterns within or between AQ(G)P groups. Here, we aim to fill this gap by analyzing the complete non-redundant set of 20 currently available high-resolution AQ(G)P structures from human and mammals, bacteria, plants and fungi available in the PDB database, with resolution between 0.88 and 3.7 Å (Table 1). The multitude of AQ(G)P structures offers the unique possibility to study the structural diversity of a class/family of proteins, providing an extensive overview of the differences in scaffold and amino acid distribution, with a strong focus on protomer/tetramer stability, pore geometry and channel functionality. This examination brings us one step further in the desire to understand the molecular architecture/construction plan of aquaporins and membrane channels in general. In this respect, AQ(G)Ps can be seen as a model case of how to conduct similar analysis on other protein families, i.e. urea channels (96, 97), in the future. Moreover, our results presented herein lay the ground to predict the potential structural and functional effects of disease-causing mutations in AQ(G)Ps.

**Table 1.**
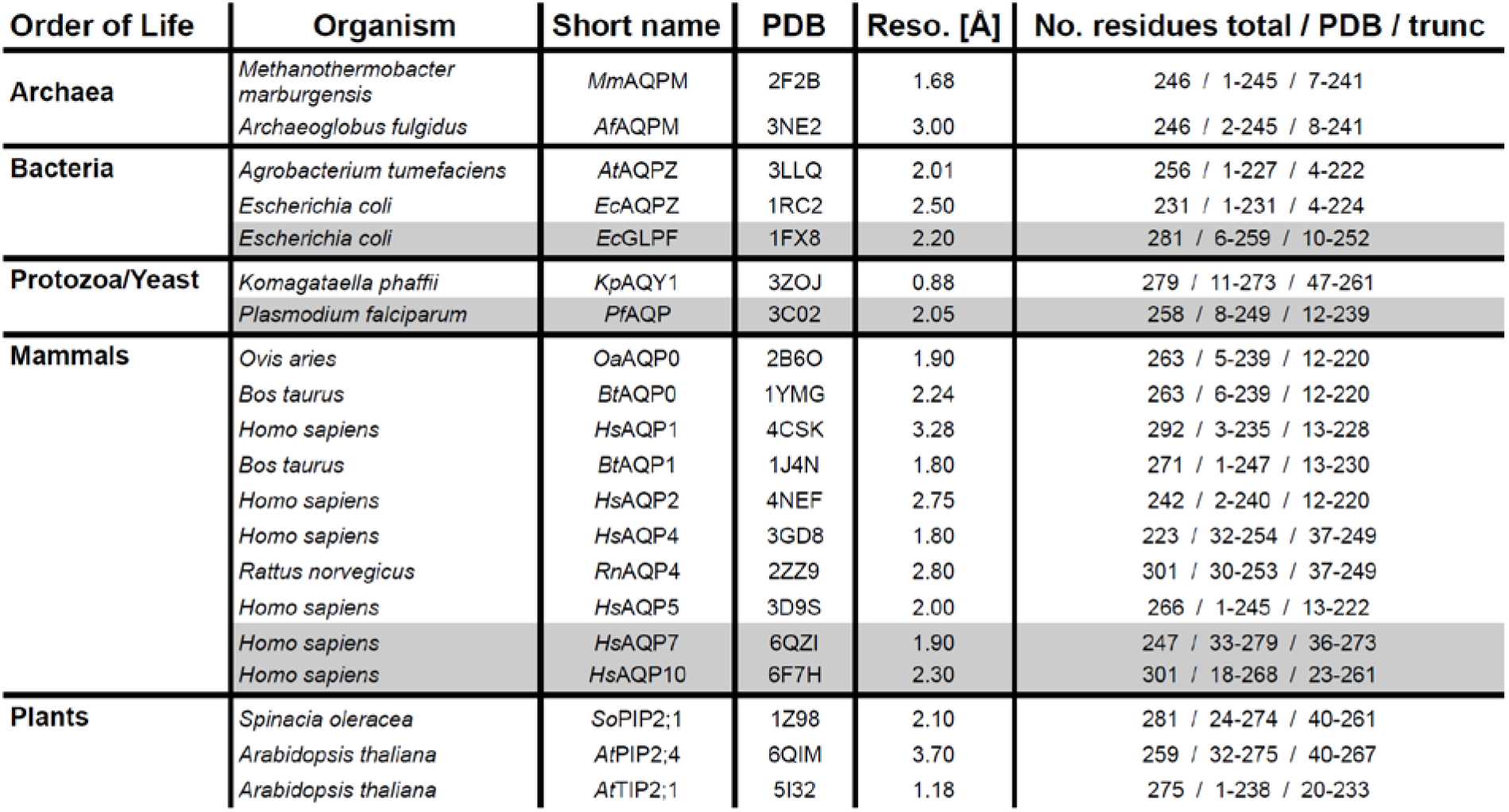
List of 20 aquaglyceroporin structures used in this study. A non-redundant list of 16 AQP and 4 AQGP (grey) structures with resolution better than 3.7 Å, excluding low resolution and lower resolved redundant structures as well as mutant proteins. MmAQPM (41), AfAQPM, AtAQPZ, EcAQPZ (42), EcGLPF (43), KpAQY1 (44), PfAQP (45), OaAQP0 (46), BtAQP0 (47), HsAQP1 (48), BtAQP1 (49), HsAQP2 (50), HsAQP4 (51), RnAQP4 (52), HsAQP5 (53), HsAQP7 (54), HsAQP10 (55), SoPIP2;1 (56), AtPIP2;4 (57), AtTIP2;1 (58). A full list of currently 60 AQ(G)P structure entries available in the PDB databank can be found in the supplementary information (**Table S1**).

## 2. Results & Discussion

### 2.1. AQ(G)Ps exhibit a common scaffold

AQ(G)Ps are homo-tetrameric membrane channels, in which the functional pores reside within each of the four protomers. As an exception, formation of PIP hetero-tetramers in plants (98–100) may represent a novel mechanism to adjust water transport across the plasma membrane (101). Interestingly, assemblies of multiple tetramers into orthogonal arrays of particles have been found for an isoform of AQP4 (102–104), with suggested but unproven functional implications on water permeability (105, 106), cell-cell adhesion (51, 102, 107), and AQP4 polarization to astrocyte end-feet (108, 109). Transport of ions and gas molecules through the central fifth pore is debated, with missing proof of biological significance (75, 110, 111). Generally, each AQ(G)P protomer consists of 6 transmembrane helices (H) and two half-helices (HH), connected by loops (L). The two half-helices meet at the membrane center and form one out of two selectivity filters within the single-file pore. The opposite dipole moments of the half helices (38–40) with the highly conserved dual NPA motives at their endings flip water molecules in the middle of the single-file region to disrupt proton permeation via the Grotthuss mechanism (112). The ar/R selectivity filter accommodates the highly conserved positively charged arginine, the second electrostatic barrier to suppress proton permeation through AQ(G)Ps, and two aromatic residues, together constituting a major checkpoint for solute discrimination (38) in the narrowest part of the pore. These fundamental AQ(G)P features, the 3-dimensional arrangement of α-helices in the protomer, as well as the oligomeric assembly are depicted in **Figure 2A-C**, with bacterial *Ec*AQPZ serving as the archetypical template due to its minimal sequence length, high structural stability, and selectivity for water. The AQ(G)P topology and fold illustrate that the N- and C-termini, as well as loops L2, L3, and L5 are oriented towards the cytoplasm whereas loops L1, L4, L6, and L7 are oriented towards the periplasm. Helices H6, HH1, HH2, and H3 are facing towards the lipid membrane and H1, H2, H5, and H4 form most of the protein-protein interface.

**Figure 2.**
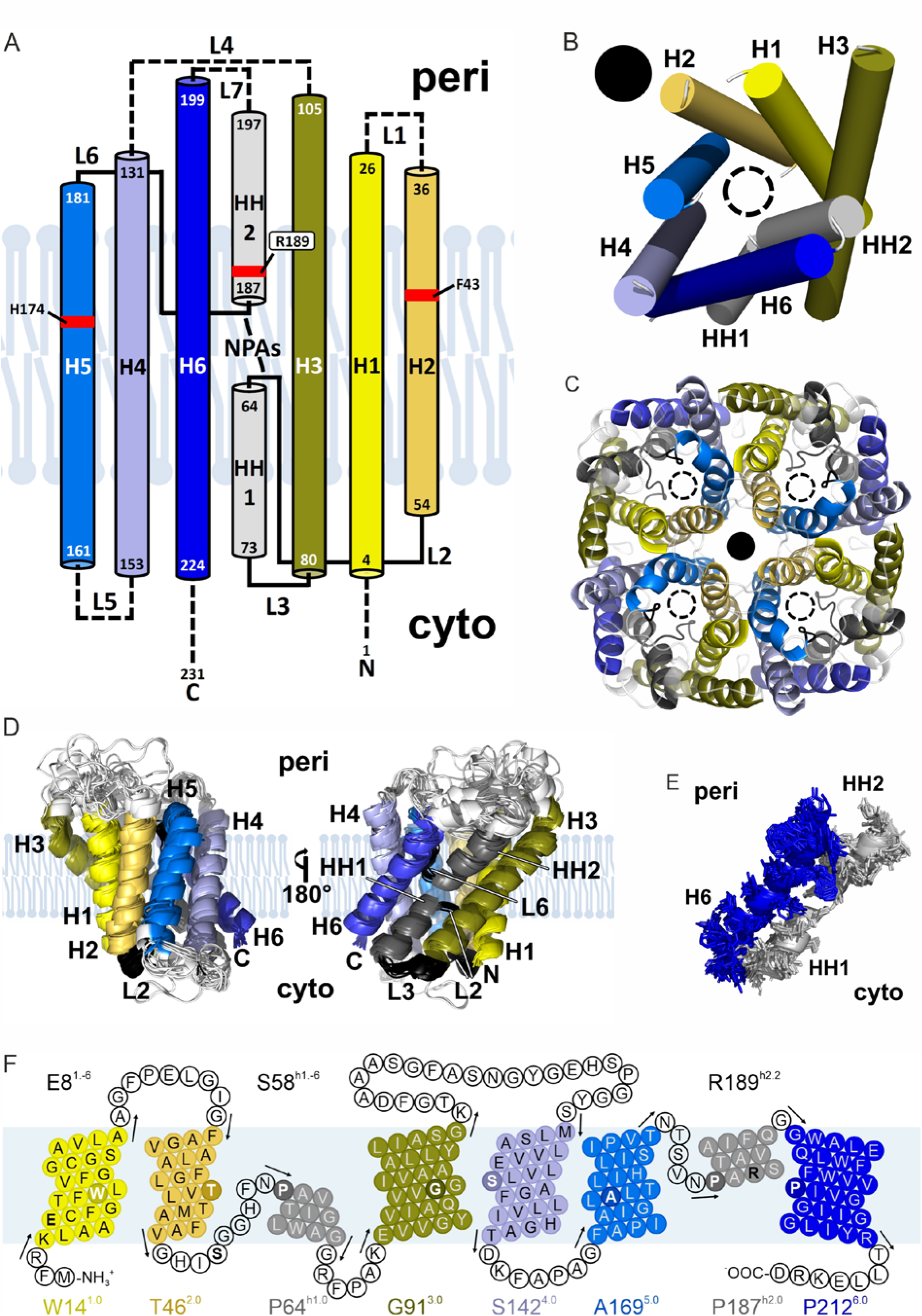
The conserved AQ(G)P scaffold. **(A)** Topology map of EcAQPZ. The NPA regions as well as residues of the ar/R selectivity filter (red boxes) are highlighted. Variable loops and termini are depicted with dashed lines. (**B**) Transmembrane helix arrangement of the EcAQPZ protomer (periplasmic view) showing the color-coding of the transmembrane helices as used throughout the manuscript. The water pore is indicated as a black dashed lined circle and the central pore in the tetrameric AQP as a black filled circle. (**C**) Tetrameric assembly of AQ(G)P protomers (periplasmic view of the EcAQPZ structure). Pore positions are indicated as in (C). **(D)** Alignment of the 20 nonredundant AQ(G)P structures using PyMOL reveals a very conserved AQ(G)P scaffold. Loop regions in white are more mobile and variable. The rest of the structure after omitting these mobile loop regions is called “core” throughout the manuscript. **(E)** Representative structural conservation along single amino acid positions of H6, HH1, and HH2. **(F)** Generalized numbering scheme illustrated on a snake-plot representation of EcAQPZ. White bold letters in darker circles represent the residue at the center of the membrane of the corresponding helix, serving as a reference residue in the numbering scheme and are listed at the bottom of each helix. Black bold letters illustrate three examples of highly-conserved residues, i.e. Glu8^1.-6^ (**2**), Ser58^h1.-6^ (**2**), R189^h2.2^ (**\***).

Alignment of all 20 non-redundant AQ(G)P structures revealed a perfectly conserved transmembrane protein scaffold **(Figure 2D**) with variations in the loops, N- and C-termini. Therefore, we defined a common scaffold of all AQ(G)P structures shown in **Figure 2D** for further structural considerations. Regions showing some variability in transmembrane helix and loop length are excluded from most of the structural analysis further on. The structural alignment does not only visualize the highly conserved α-helical assembly but also the perfectly structurally conserved loops L2, L3, and L6, connecting H2 with HH1, HH1 with H3, and H5 with HH2, respectively. A closer look at the protein backbone **(Figure 2E**) shows that it is possible to assign each amino acid in the common scaffold to a respective 3D position within the structure, further supporting the notion of a universal AQ(G)P fold. Moreover, the precisely conserved Cα positions of individual amino acids of all AQ(G)Ps analyzed in this work will confer confidence to expand the analysis of amino acid distributions - including their chemical properties - at secondary or tertiary structural interfaces to a multitude of AQ(G)P sequences with yet unresolved structures in the future.

### 2.2. Numbering scheme for AQ(G)Ps

The universal AQ(G)P fold with its conserved Cα positions allows us to introduce a numbering scheme for AQ(G)Ps similar to the Ballesteros-Weinstein numbering of G protein-coupled receptors (GPCRs) which was already introduced in 1995 (113). This will provide a framework to relate site-specific properties to the sequences of structurally yet unresolved AQ(G)Ps facilitating structural and functional comparisons and predictions of e.g. disease-causing mutations in mammalian AQ(G)Ps. The numbering scheme is informative of the amino acid (AA) present at that position, the real AA number in a particular AQ(G)P, and the relative position of AAs. All these AA position-specific pieces of information are condensed in identifiers, derived as follows: The AA is labeled by a standard one or three letter code and its real AA number, then the AA identifier in superscript starts with the helix number (1–6), e.g., 1, for helix H1. To avoid confusion HH1 is denoted as, h1, and HH2 as, h2. This helix identifier is separated by a point from the position relative to a reference residue which was chosen to be the residue in the center of the lipid bilayer of the corresponding helix (see **Figure 2F**). That reference residue is assigned the number 0. For example, in H1 it is a tryptophan in position 14 in *Ec*AQPZ, whose identifier would be 1.0, i.e., Trp14^1.0^. A glycine residue located five AA after Trp14^1.0^ will be Gly19^1.5^, the glutamate acid residue six positions earlier in the sequence is Glu8^1.-6^. This general numbering scheme is illustrated in **Figure 2F** on the snake-plot representation of *Ec*AQPZ and used throughout the manuscript to relate AA positions of different AQ(G)Ps to each other. Mutations are identified in this numbering scheme in the usual manner, with the wild-type residue followed by the mutant AA. For example, E8K^1.-6^ defines the Lys mutation of the wild-type Glu in helix 1, residue 8 in the *Ec*AQPZ sequence, in the position 1.-6. The identification scheme allows for a systematic comparison of mutations done in different AQ(G)Ps at the same location. Moreover, the assigning of the reference residue to the mid of the bilayer allows for rapid comparison of the helix length on the periplasmic or cytoplasmic side. **Figure 2F** illustrates the selected reference AAs in each helix with the corresponding identifiers according to *Ec*AQPZ as well as three representative identifiers of conserved AAs. According to our numbering scheme, AAs in loops can principally have two identifiers, e.g., Ser58^h1.-6^ = Ser58^2.12^. However, we suggest using the identifier related to the next C-terminal helix, e.g. Ser58^h1.-6^ in our example, although both define uniquely the same position. This has the advantage that L2 and L6 are affiliated with the identifiers h1 and h2 of the neighboring half helices HH1 and HH2, respectively. However, it might also be reasonable to number N-terminal residues of L1, L4, L5, and L7 according to their adjacent N-terminal helix, as this emphasizes the structural vicinity and conservation. Clearly, AQP identifiers depict a unique position within the AQ(G)P scaffold if the residue is located within the AQ(G)P core only. Identifiers for AAs located in L1, L4, L6, L7, the N-terminus, and the C-terminus are only relative in nature, as the exact position may vary among AQ(G)Ps due to differing lengths and folds of the corresponding structural element.

### 2.3. Structural features & basic differences of AQ(G)Ps

Despite a common scaffold, AQ(G)Ps are not identical, showing distinct differences. A sequence alignment (**Figure S1**) illustrates that all AQ(G)Ps exhibit NPA motives at the ends of HH1 and HH2 except *Pf*AQP and *Hs*AQP7, which possess NLA and NAA motifs at the end of HH1, respectively, and NPS instead of NPA at the end of HH2. While Arg189^h2.2^ (in *Ec*AQPZ), as part of the three aromatic residues in the ar/R filter, is perfectly conserved among all AQ(G)Ps, Phe43^2.-3^ and His174^5.5^ slightly vary. Phe43^2.-3^ in *Ec*AQPZ corresponds to a Trp in *Ec*GLPF and *Pf*AQP and a His in *At*Tip2;1. In *Hs*AQP10, Gly61^2.-3^ at this position provides space for Phe58^2.-6^, reaching into the same space from one α-helical turn above (three residues earlier, towards the N-terminus and periplasm). The second aromatic residue, His174^5.5^ (*Ec*AQPZ) is replaced by an Ile in *At*Tip2; 1 and AQPMs or by Gly in all AQGPs. In the glycerol conducting aquaporins *Ec*GLPF, *Pf*AQP, *Hs*AQP7, and *Hs*AQP10, His174^5.5^ is functionally substituted by Phe, Phe, Tyr and Ile, located in L6, 9 residues further to the C-terminus (identifier h2.-4), respectively. This illustrates quite plainly the difference between AQGPs, AQPs, AQPMs and of the ammonia permeable *At*Tip2; 1 concerning their solute selectivity. Our classification of AQ(G)Ps into these four groups for further analysis throughout the manuscript is confirmed by the sequence similarity map (**Figure S2**) and the polygenetic tree of the 20 candidate sequences (**Figure S3**). Overall, the sequence similarity of truncated AQ(G)Ps varies between 58.3% for *At*PIP2.4 and *Hs*AQP10 and 86.0% for *Hs*AQP2 and *Hs*AQP5. Similarly, denominated AQPs from different organisms vary between 86.5% and 86.9% for AQPZ and AQPM to 98.9% and 98.1% for AQP0 and AQP4, respectively.

Focusing on the length of helices and loops, the following observations were made. While the TM helix length of HH1 is constant in all investigated AQ(G)P structures, HH2 is longer for AQGPs and AQPMs compared to AQPs (**Figure S4**). No clear pattern can be observed regarding the length of H2, H4, and H5. H6 is shorter in most mammalian AQPs except *Hs*AQP7 and H3 is strikingly longer in AQGPs towards the periplasmic side. The length of H1 is generally most diverse, being longest in plant AQPs and *Kp*AQY1. The lengths of L2, L3, and L6 are perfectly conserved, showing 9, 6, and 5 residues, respectively, whereas L5 is longer in plant PIPs and slightly longer in AQGPs (**Figure S5**). The length of L1 is most diverse among all loops, with AQPMs having by far the longest loop with 25 amino acids compared to only 6 residues in both AQP0. L4 is much longer in AQGPs (*Ec*GLPF, *Hs*AQP7, *Hs*AQP10), slightly longer in *At*AQPZ, *Ec*AQPZ and *Pf*AQP, and shortest in *At*TIP2;1 and *Kp*AQY1. L7 is longest in AQGPs and slightly elongated in AQPMs and PIPs. Despite some AQ(G)P specific variations in helix and loop length, there are obvious differences between AQGPs and AQPs in the length of H3 and HH2 as well as the corresponding periplasmic loops L4 and L7, connecting H3-H4 and HH2-H6, respectively. Both the loop and N- and C-termini diversity suggests regulatory functions via the interaction with peripheral proteins (94). In this respect, archaeal and bacterial AQ(G)Ps as well as *Hs*AQP7 exhibit the shortest termini by far (**Figure S5**). Having a closer look at the loops apparent polarity confirms the positive-inside rule (114), with basic residues being markedly more abundant on the cytoplasmic side (**Figure S6 – S8**). Nevertheless, the extend and distribution of surface electrostatics varies greatly between the 20 AQ(G)P structures (**Figure S8**), in compliance with the possibility to serve as binding platform for peripheral proteins.

### 2.4. Disparities in pore geometry and radius determine solute permeability

Classical AQPs exhibit a long, narrow single-file region with the narrowest part formed by the ar/R selectivity filter. This can be seen by a channel radius of 0.8-1.8 Å for the blue colored exemplary pore radii in **Figure 3A** and the long narrow (green surface) pore surface in **Figure 3B** for *Ec*AQPZ. In case of *Kp*AQY1, Tyr31^1.-26^, part of a putative gating mechanism, occludes the pore on the cytoplasmic side of the channel visualized in **Figure S9**. Mutational studies in combination with molecular dynamics simulations imply that gating by Tyr31^1.-26^ may be regulated by a combination of phosphorylation and mechanosensitivity in *Kp*AQY1 (115). AQGPs (red lines in **Figure 3A**) show a much larger pore radius throughout most of the pore, as is apparent for the respective pore surface representations of *Ec*GLPF and *Hs*AQP10 in **Figure 3B**. Interestingly, AQPMs show an intermediate pore radius, with a broad pore geometry at the cytoplasmic side similar to AQGPs, yet a narrow ar/R filter at the periplasmic end of the channel, similar to AQPs. This mixed shape of the pore correlates well with AQPMs solute selectivity and the evolutionary analysis (shown in **Figure S3**). In detail, (i) AQPM has been observed to transport glycerol, albeit at lower rates as compared to *Ec*GLPF (116). (ii) Moreover, a glycerol molecule was bound inside the pore in the crystal structure of *Mm*AQPM (41). (iii) Finally, the evolutionary analysis revealed that AQPMs are an intermediate between AQPs and AQGPs (7, 117, 118). However, the biological relevance of the observed increased permeability to larger neutral solutes is questionable in case of the primary host of *Mm*AQPM, *Methanothermobacter marburgensis*, as it relies on CO_2_ as its sole carbon source (116), and thus does not require a dedicated glycerol facilitator for survival. The hyperthermophilic sulphate-reducing archaeon *Archaeoglobus fulgidus* additionally encodes a glycerol facilitator *Af*GLPF (119), utilizing the glycerol derivative diglycerol phosphate as an osmolyte under high-salt conditions (120, 121).

**Figure 3.**
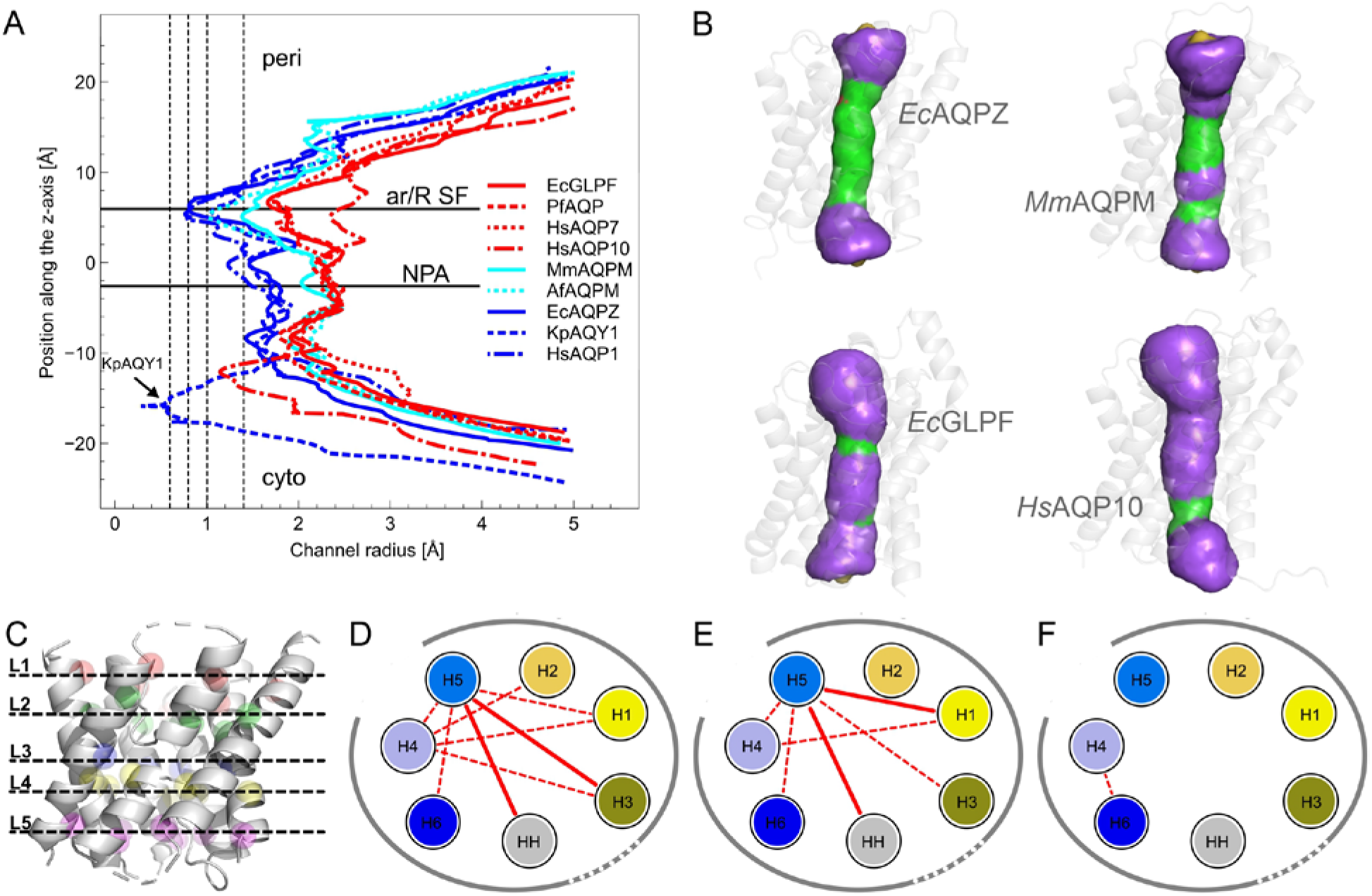
Pore geometries and scaffold differences of AQ(G)Ps. **(A)** Pore radii as estimated with the program HOLE (122, 123). AQGPs are indicated in red, 3 representative AQPs are labeled in blue and the two AQPM structures in cyan. The vertical lines guiding the eye highlight pore radii of 0.6, 0.8, 1.0 and 1.4 Å. **(B)** Exemplary pore surfaces of EcAQPZ, MmAQPM, EcGLPF, and HsAQP10 generated by HOLE (122, 123). Protomer structures are depicted in a transparent side view (periplasm up / cytoplasm down). The color scheme according to the program HOLE is the following: Green areas represent pore radii suitable for a single water molecule only, red colored pore surfaces are too narrow to let water molecules pass, and purple regions can accommodate more than one water molecule per cross section. Pore radii and pore geometries of our full set of non-redundant AQP structures can be found in the SI (**Figure S9, Figure S10**). **(C)** Positions of the 5 layers (shown for EcAQPZ) used to determine helix-helix distances along the channel (shown in **Figure S14**). **(D)** Averaged helix-helix distance differences of AQGPs to AQPs, **(E)** AQGPs to AQPMs and **(F)** AQPMs to AQPs. Increase of distance by 0.5 to 1 Å is shown by a dashed red line and above 1 Å by a solid red line.

### 2.5. Solute-specialized aquaporins differ subtly in the AQ(G)P scaffold structure

AQGPs exhibit a wider pore geometry than AQPs in order to accommodate and conduct neutral solutes like glycerol **(Figure 3A**). However, as mutational studies of *Ec*AQPZ engineered with three signature amino acids of *Ec*GLPF (F43W^2.-3^/H174G^5.5^/T183F^h2.-4^) suggest (62), the wider pore geometry and solute selectivity of AQGPs is not solely achieved by a different set of pore lining residues. This proposes that in addition to pore lining residues, different relative positions of one or more α-helices within each protomer to each other could play an equally important role in shaping the NPA region and the ar/R selectivity filter. To locate these subtle structural differences, we first analyzed the RMSD of all AQP structures to each other (**Figure S11**). The RMSD plot roughly confirms the polygenetic tree (**Figure S3**) and the sequence similarity plot (**Figure S2**), clearly grouping AQGPs or mammalian AQPs together. Interestingly, the analysis revealed that the mammalian AQP1 is very much different from the rest of mammalian AQPs, except for AQP0 and AQY1, as well as from bacterial and archaeal AQPs. Plant PIPs are different from *At*TIP2;1 and show a high RMSD when compared to basically all other AQPs. The analysis also revealed distinct and unexpected differences between similar proteins from different organisms, as for AQPMs. However, general differences between AQPs and AQGPs could not be established at this general level.

For a more detailed analysis, we next concentrated on the variance of single Cα positions. A comparison between all 20 structures (**Figure S12**), with *Ec*AQPZ as a reference, revealed that the periplasmic side is generally more diverse as compared to the core and the cytoplasmic side. Helices H2 and H5 in AQGPs deviate from the rest of the AQPs. *At*TIP2;1 differs on the cytoplasmic side of H2 similar to the AQGPs. AQPMs and PIPs are rather similar to AQPs as compared to AQGPs. On the other hand, the position of the NPA motive as well as the vicinal amino acids and the aromatic arginine containing selectivity filter are universally conserved.

In order to gain information about the relative directional deviation of single α-helices to each other, we had a closer look on helix-helix distances in 5 layers along the membrane normal (**Figure S13**). Thereby, layer position 1 is located at the periplasmic side, layer position 3 in the middle of the membrane and layer position 5 at the cytoplasmic side **(Figure 3C**). To find differences between AQPs, AQGPs and AQPMs, we further averaged the respective helix-helix distances into these three groups (**Figure S14**). Visualization of the results reveals subtle differences in the relative α-helix positions within AQGPs as compared to AQPs for each of the five layers (**Figure S15A**). Whereas the cytoplasmic vestibule (layer 5) seems to be generally wider, in the center of the membrane, at the height of the periplasmic NPA motive (layer 3), the major difference is an increased distance between H5 and helices H3, H4, as well as H6 by more than 1 Å. An average over all 5 layers (**Figure S16**) illustrates the major differences in the relative backbone arrangements between AQPs and AQGPs. That is, H5 is shifted towards the central pore of the tetramer, leading to an increase distance to mainly H3 and HH1. Moreover, the distance of H4 to H1, H2, H3 and H5 is increased. Together with the increased distance between H4 and H5, these results point to a larger spread of the AQGP helices compared to AQPs. **(Figure 3D**). The increased distance c between H4 and H5 could be further verified by measuring its center of mass distances (**Figure S17**). Furthermore, center of mass distances of the inter-protomer distance between contacting helices H2:H5, single protomers, and the corner half-helices HH (shown in **Figure S17**) revealed both that the distance between protomers of AQGPs is larger by 1.1 Å, and the overall extension of tetrameric AQGPs is larger by 2.6 Å. The differences in the interhelix distances between AQGPs and AQPMs are similar to those of AQGPs and AQPs (**Figure 3E**). AQPMs and AQPs on the other hand are rather similar in their averaged helix-helix distances (**Figure 3F**).

However, more detailed view on the individual layers in **Figure S15** shows that the cytoplasmic vestibule of AQPMs is slightly tighter, with H1 in closer contact to H2, and the periplasmic vestibule slightly wider, with H3 oriented outwards the channel center.

The overall increased pore geometry as well as a wider scaffold of AQGPs compared to AQPs strengthens the notion that the selectivity filter is also shaped by the backbone (distances of helices to each other) and not solely by the corresponding side chains. As AQPMs exhibit a similar scaffold as compared to AQPs **(Figure 3F**) but a comparable pore geometry to AQGPs with low glycerol permeability **(Figure 3A**), we speculate that it is possible to reach glycerol selectivity with an AQP backbone, yet at low permeability. High glycerol permeabilities require an adaptation of the relative helix positions to each other.

### 2.6. Water is guided through the AQP pore along a line of H-bond forming residues

The mobility of pore water in AQ(G)Ps was suggested to be governed by the number of H-bonds, N_H_, water molecules may form with pore lining residues (4). The mobility increases in a logarithmic dependence with higher N_H_, in line with the multiplicity of binding options at higher N_H_ densities. Except for the ar/R selectivity filter with a multitude of H-bond forming options for single-file water molecules, water is guided through the cytoplasmic half of the channel by forming H-bonds with residues on one side of the channel only, likely in order to stabilize the orientation of the water molecules in the pore and to allow for rapid water flow (**Figure S18**). As depicted in **Figure 4** and **Figure S19**, most of these interactions occur with backbone oxygens (residues labeled a, b, c, d, i, j, k, l, and m) as weak H-bond acceptors, whereas only 5 residues form H-bonds to water via their side chains (labeled e, f, g, h, m). Interestingly all five of those residues are involved in the formation of the selectivity filters. Two out of these residues are the conserved Asn of the two NPA motives. The remaining three residues belong to the ar/R selectivity filter: Arg^h2.2^ in position f, a prevalently conserved His^5.5^ (His174^5.5^ in *Ec*AQPZ, position g), and position m. The lattest position is filled with a Phe^2.-3^ in most of the AQ(G)P structures except for *Ec*GLPF, *Pf*AQP, and *At*TIP2; 1, which exhibit Trp^2.-3^, Trp^2.-3^ and His^2.-3^, respectively. Only Trp^2.-3^ and His^2.-3^ are able to contribute in H-bond formation by their side chains. Overall, this greatly limits the variability in the number of H-bonds single-file water molecules can form with pore lining residues in position g and m. Generally, nitrogens are thought to build stronger H-bonds than oxygen atoms (124), with the strength of H-bonds formed with water having a pronounced effect on the H-bond dynamics (125) and inducing a moderate slowdown in the water H-bond exchange dynamics due to an excluded volume effect (125), similar to that of hydrophobic groups. Such H-bonds are in AQ(G)Ps formed with side chains of the five above mentioned amino acids **(Figure 4A** labels e, f, g, h, m). In *Ec*AQPZ, the side chain of Asn182^h2.-5^ (which is a Gly in all other 19 structures) is also capable of H-bonds formation. All other H-bond donor groups are backbone oxygens, likely designed by nature for increased water permeability. Detailed analysis of the average number of H-bonds of pore lining residues with pore water molecules in our MD simulations revealed interesting differences (**Figure S20**). In detail, on average 16.0, 13.6, 19.5 and 18.6 H-bonds were observed for *Ec*AQPZ, *Bt*AQP1, *Hs*AQP4 and *Ec*GLPF, respectively. As the rest of the narrow pore is decorated by hydrophobic residues only, we speculate whether substitution of those hydrophobic residues for AAs capable of forming H-bonds would disturb rapid sliding of water molecules through the narrow pore as well as interfere with high permeabilities of AQ(G)Ps.

**Figure 4.**
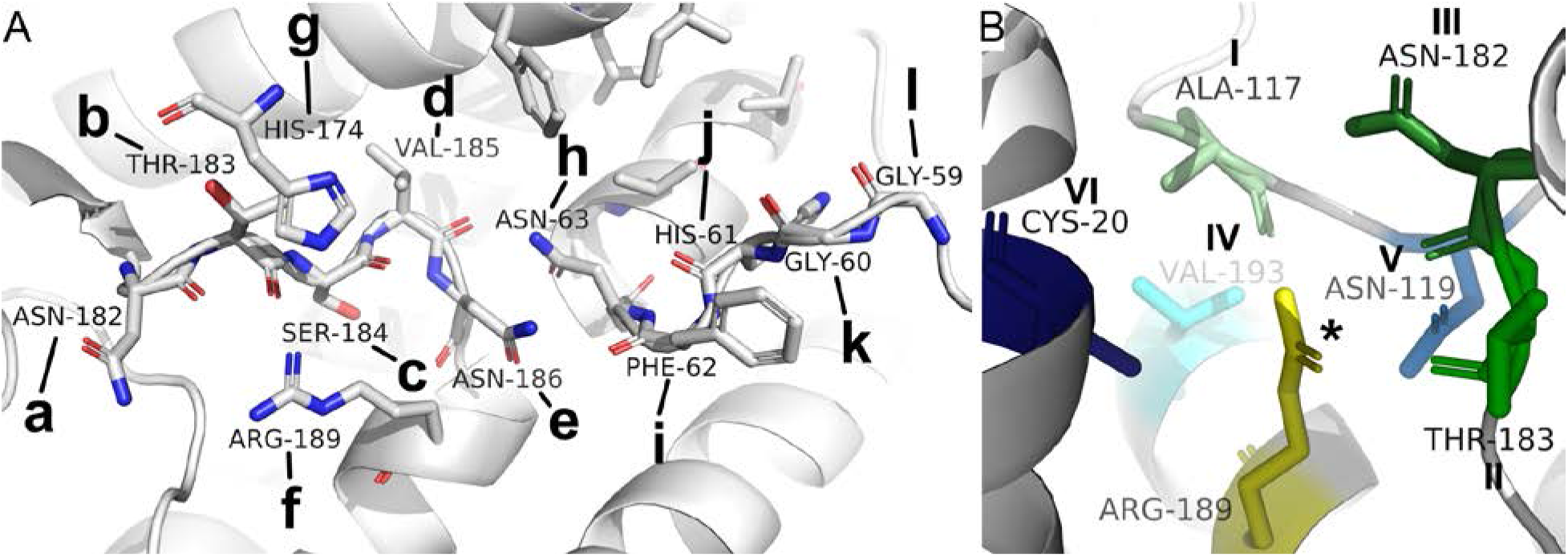
Line of H-bond donors and acceptors guide water through the AQ(G)P pore. (A) Residues in our template EcAQPZ potentially forming H-bonds to passing water molecules according to the sub-Ångström structure of KpAQY1, which for the first time directly visualized H-bonds of crystalized single-file water molecules in the pore with pore lining residues. Through visual inspection of potentially solvent exposed residues in the single-file pore of EcAQPZ, we additionally included Phe62^h1.-2^ and Val185^h2.-2^. Indeed, our MD simulations have shown that residues in these two positions are not only capable to form H-bonds with pore waters in EcAQPZ, yet also in HsAQP1, HsAQP4 and EcGLPF (see **Figure S20**). Position m (not shown) is only relevant in EcGLPF, AtTIP2;1 and PfAQP as all other 17 AQ(G)Ps exhibit a Phe^2.-3^ in this position (**Figures S19**). Residues are labeled from left (periplasm) to right (cytoplasm). For a cytoplasmic and periplasmic view, see **Figure S18**. (**B**) Hydrogen bonding network stabilizing the Arg^h2.2^ side chain in the ar/R filter. Potential interaction partners observed in the 20 nonredundant AQ(G)P structures are highlighted in the EcAQPZ structure and colored based on the classification of all individual positions (**Figure S21**). H-bonding with Arg^h2.2^ itself and amino acids I, II, and III occurs with the backbone, whereas amino acids IV, V, and VI interact via their side chains.

### 2.7. AQ(G)P gating

Even though AQ(G)Ps are commonly seen as constantly open passive facilitators of water and other neutral solutes, there is a plethora of mechanisms reported influencing their transport capabilities. Water passage can be modulated by movement of side chains of pore lining residues into the pore (56, 87, 88, 90, 126) or even by large scale rearrangements of structural elements (109). A similar process involving two periplasmic loops is also thought to gate the proton gated inner membrane urea channel of *Helicobactor pylori*, *Hp*UreI (127). In AQ(G)Ps, structurally caused permeability modulations have been proposed to be triggered by pH (89), divalent cation binding (128), phosphorylation (129), mechanical stress (115) and protein binding (130) and can act at different positions along the pore.

#### 2.7.1. Arg in the ar/R filter region

Numerous indications from *in silico* (61, 131–134) and *in vitro* (42, 135, 136) studies point to a conserved and highly flexible Arg^h2.2^ in the ar/R selectivity filter able to block water passage through the single-file water pore of AQPs. This could provide AQPs with a putative transmembrane voltage or lipid asymmetry dependent regulation mechanism. However, gating by the conserved Arg^h2.2^ is still under debate (137), with a clear experimental proof missing. Herein, we analysed the hydrogen bond network, stabilizing the position of the side chain of Arg^h2.2^ in the channel pore in the 20 high-resolution structures and of four selected targets also by MD simulations, in order to gain a view on its dynamics. In our structural analysis, six neighboring residues were determined which form at least one contact with Arg^h2.2^ in one of the 20 structures (**Figure S21**). For illustration, those seven residues (including Arg^h2.2^) were mapped onto the *Ec*AQPZ structure **(Figure 4B**). It is important to note that no AQ(G)P structure showed interaction with all of those 7 residues at once. The nitrogen atoms on the Arg189^h2.2^ side chain may form H-bonds to the backbone oxygens of following residues: Arg^h2.2^ itself (labeled *), the position of Ala117^4.-25^ (I) in L4 and the positions of Asn182^h2.-5^ (III) and Thr183^h2.-4^ (II) in L6. Furthermore, we discovered Ser^h2.6^ in the position of Val193^h2.6^ (IV) in HH2, Asn^4.-23^ in position of Asn119^4.-23^ (V) in L4 and Glu^1.6^ in the position of Cys20^1.6^ (VI) in H1 as potential side chain H-bond interaction sites with the respective Arg^h2.2^ in certain AQ(G)Ps. Interestingly, the presence of Asn^4.-23^ does not necessarily imply formation of an H-bond to Arg^h2.2^. Fascinatingly, the overall amount of H-bonds stabilizing the respective Arg^h2.2^ varies widely, with a minimum of one H-bond in *At*Tip2;1, two H-bonds in *Ec*GLPF, *Hs*AQP7 and *Hs*AQP10, five H-bonds in *Pf*AQP and *At*AQPZ, and a maximum of six potential H-bonds counted in *Hs*AQP5.

To verify the structurally found differences in H-bond stabilization of the conserved Arg^h2.2^ and to thereby be able to explain and predict potential Arg^h2.2^ gating *in vivo*, we conducted MD simulations of selected AQ(G)Ps. Indeed, all atom MD simulations of *Bt*AQP1, *Hs*AQP4, *Ec*AQPZ, and *Ec*GLPF confirmed the Arg^h2.2^ H-bond interaction partners found in our structural analysis (**Figure S22**), with the exception that Arg189^h2.2^ in *Ec*AQPZ does also form an H-bond with itself for about 50% of the simulation time and the H-bond to Thr183^h2.-4^ is almost never observed in MD simulations. Also, the probability for Arg197^h2.2^ to form a H-bond to Ser201^h2.6^ in *Bt*AQP1 amounted to only 20%. In addition, it got obvious that the H-bond probability between individual AQP protomers was rather homogeneous for *Bt*AQP1 and *Ec*AQPZ, indicating a relatively stable conformation, while the H-bond network in *Hs*AQP4 was different in chain D compared to the other protomers, hinting at a less stable H-bond network and higher Arg^h2.2^ flexibility. Strikingly, in *Ec*GLPF, the situation was completely different, exhibiting a very unstable and disperse H-bond network of neighboring residues with Arg^h2.2^. This finding is underpinned with on average 0.83±0.12, 1.84±0.11, 2.31±0.04, and 2.56±0.04 H-bonds with neighboring aminoacids in *Ec*GLPF, *Hs*AQP4, *Ec*AQPZ, and *Bt*AQP1, respectively. We further visualized the flexibility of Arg^h2.2^ in *Ec*GLPF and chain D of *Hs*AQP4 by plotting the angle of the respective arginines in the pore. As it can be seen in **Figure S23**, Arg^h2.2^ exhibits indeed larger flexibility. Moreover, the orientation of Arg^h2.2^ in the pore modulates water passage though the respective pores (**Figure S24**). While Arg^h2.2^ leads to an open pore at a dihedral angle of around 300° and 100°, the pore is closed at an intermediate dihedral angle of 180° (Figure 5, subset i). Hence, we conclude that the flexibility of Arg^h2.2^ in the pore varies significantly between AQPs, having a severe impact on the rate of water passage through the respective channel. The number of H-bonds found in AQP structures stabilizing Arg^h2.2^ may serve to predict this tendency *in silico* and potentially also *in vivo*. We speculate that Arg^h2.2^ in *Hs*AQP7, *Hs*AQP10 and *At*TIP2;1 are highly flexible as in the case of *Ec*GLPF, but Arg^h2.2^ in AQPMs, *So*PIP2;1, and *Rn*AQP4 may be similarly stable as compared to *Hs*AQP4. *Hs*AQP5, *Kp*AQY1, *At*AQPZ, and *Pf*AQP seem to have an even more refined H-bond network comprising one or two additional H-bonding partners.

**Figure 5.**
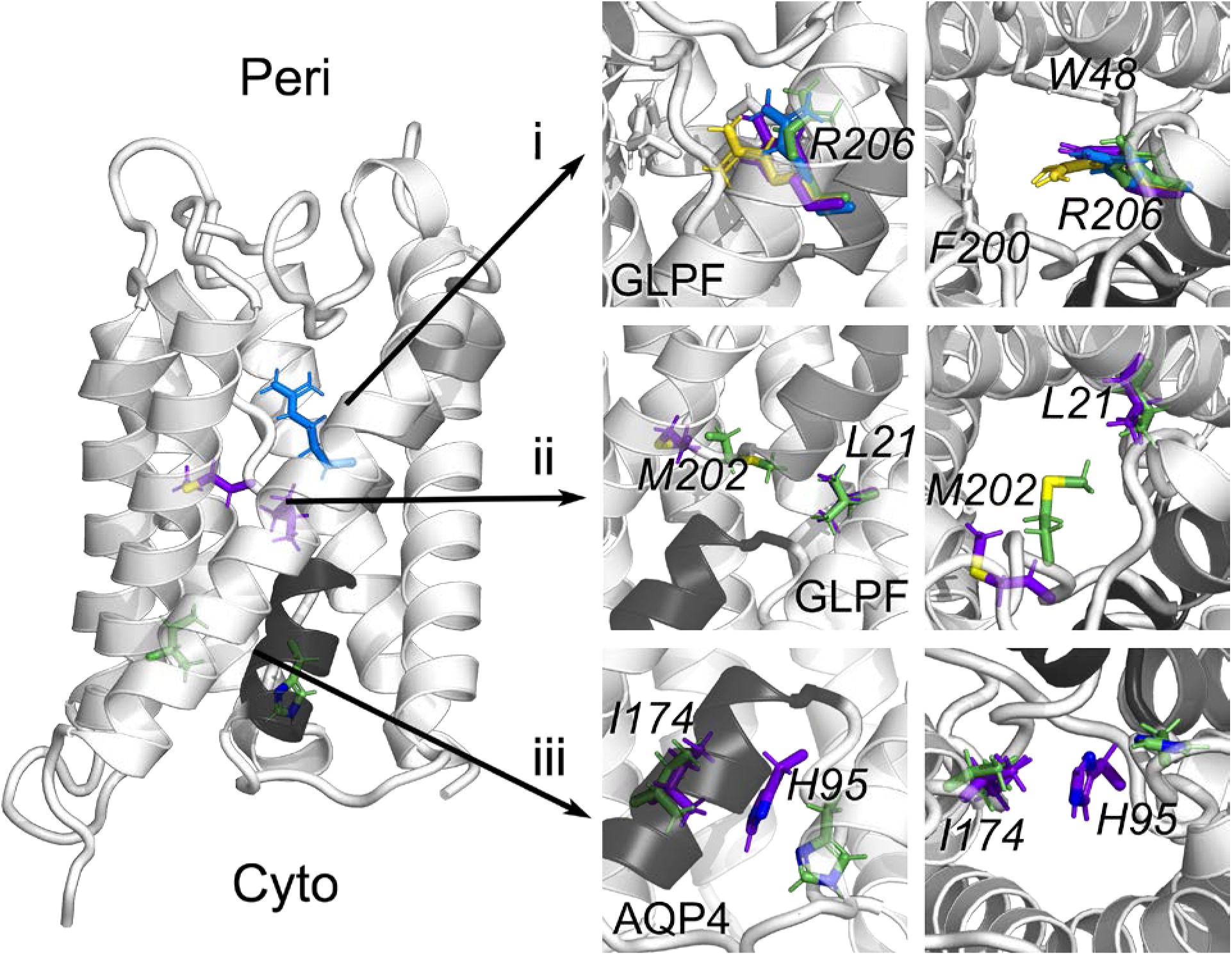
AQ(G)P gating in MD simulations. MD simulations of BtAQP1, HsAQP4, EcAQPZ, and EcGLPF have revealed three possible dynamic gating sites by residues moving into the pore lumen. Gating site **(i)** is the conserved Arg^h2.2^ in the ar/R selectivity filter. Arg^h2.2^ can occupy four different positions, described here by the dihedral angle Cα-Cβ-Cγ-Cζ. The angle of 320° (colored green) represents an open state in the position of Arg^h2.2^, similar to that observed in most crystal structures. Arg^h2.2^ in BtAQP1 and EcAQPZ only rarely leaves this position. The angle of ~270° (colored blue) represents another open state, however, with the guanidyl group slightly more protruding to the pore and thus slowing down the water passage. While the pore is blocked by Arg^h2.2^ at the angle of ~200° (colored yellow), further reduction of the dihedral angle to ~100° (colored purple) results in pore opening again. The second restriction site **(ii)** is characterized by large scale movement of Met^h2.-2^ into the pore lumen, here shown for Met202^h2.-2^ in EcGLPF and quantified by measuring the minimum distance of Met202^h2.-2^ to Leu21^1.1^. At short distances (<0.4 nm) the pore is blocked (shown in green), at larger distances (colored purple) the pore is open. The third constriction site is the highly conserved His^h1.-3^ in the cytoplasmic pore lumen **(iii)**. The pore closure is visualized for HsAQP4 and measured by estimating the minimal distance between the His95^h1.-3^ and Ile174^4.7^. Small distances (shown in purple) result from flipping of the histidine into the pore and lead to its closure. At long distances (His95^h1.-3^ and Ile174^4.7^ shown as green sticks) the pore is open. The middle panels show side views on the constriction sites and the right panels show views from the periplasmic (i and ii) and cytoplasmic (iii) side into the pore.

#### 2.7.2. Dynamic constriction regions identified by MD simulations

In addition, our MD simulation unraveled two more locations where pore lining residues modulate water flux by correlating the positions of the pore lining residues with the number of water passage events through *Ec*GLPF, *Hs*AQP4, *Ec*AQPZ, and *Bt*AQP1 over time (**Figure S24**). The reduction of water permeability is thereby caused by (temporary) translation of the respective side chains into the channel pore, namely the conserved His^h1.-3^ at the cytoplasmic end of the water pore region and Met^h2.-2^ directly preceding the second NPA motif. Met^h2.-2^ is conserved in 8 out of 20 here investigated AQ(G)Ps, all other structures show hydrophobic Ile, Leu, or Val in the respective position, which anchor loop L6 well in the hydrophobic surroundings. The backbone of this residue is stabilized by a hydrogen bond with the conserved Glu^4.-4^ **(Figure 9, label 10**) in H4.

Met202^h2.-2^ positioning in the *Ec*GLPF pore, measured by a minimal distance between Met202^h2.-2^ and Leu21^1.1^, strongly modulates water passage in our simulation (**Figure 7ii**, **Figure S24**). While a distance of around 0.7 nm allocates an open pore, distances of around 0.3-0.4 nm correspond to a closed pore. Interestingly, Met212^h2.-2^ of *Hs*AQP4 in the same position does not exhibit gating behavior over the course of our simulations. The slightly reduced average minimal distance of 0.6 nm between Met212^h2.-2^ and Phe48^1.1^ in *Hs*AQP4 as compared to *Ec*GLPF is in agreement with a smaller pore radius at the methionine position (**Figure S16**, position along the pore of about 0 Å). In order to elucidate the reason for this difference, we analyzed the H-bonds stabilizing the backbone of these methionines (**Figure S23**). Indeed, the H-bonding of the conserved Glu^4.-4^ (Glu163^4.-4^ and Glu152^4.-4^ in *Hs*AQP4 and *Ec*GLPF, respectively) is much less frequent in *Ec*GLPF, compared to *Hs*AQP4. Especially, in chain B of *Ec*GLPF, exhibiting the largest closure probability by Met202^h2.-2^ (**Figure S24**), Met202^h2.-2^ is H-bonded to Glu152^4.-4^ only for 10% of the simulation time (**Figure S23**). This observation confirms our supposition that the movement of this methionine to the pore lumen is interconnected with deficient anchoring of its backbone to the rest of the protein. Recently, methionine in the hydrophobic core of the spidroin protein was shown to bestow the protein with exceptional dynamics, enabling it to adjust its shape and thus to customize its function (138). Unfortunately, the literature evidence about the role of methionine in AQ(G)Ps is scarce. Met212^h2.-2^ in *Hs*AQP4 was suggested to directly interact with His201^5.5^ in the ar/R filter and Met209^h2.-2^ in *Hs*AQP8 was suggested to be involved in protection mechanism of cells against damage by reactive oxygen species (73). For the first time, our results here point to the water-flux mediating role of Met^h2.-2^ positioned in the vicinity of the NPA and ar/R motifs inviting further experimental and simulational investigations.

Gating by the conserved His^h1.-3^ at the cytoplasmic vestibul was first suggested by Janosi et al. in 2013 for *Hs*AQP5 (139), followed by Alberga et al. in 2014 for *Hs*AQP4 (140), and soon afterwards it was shown to be further influenced by pH (141). While pH gating will be discussed in the next section, here, we concentrate on His^h1.-3^ gating in a single protonation state. Our MD simulations show that next to *Hs*AQP4 also *Bt*AQP1 and *Ec*AQPZ, unlike *Ec*GLPF, can be gated by the movement of the cytoplasmic His^h1.-3^ into the pore lumen (**Figure S24**). Our analysis of H-bond patterns between the cytoplasmic His^h1.-3^ and various interaction partners has revealed stable anchoring of *Ec*GLPFs His66^h1.-3^ by Glu14^1.-6^ and Thr72^h1.3^ (**Figure S23**).

#### 2.7.3. pH gating

Numerous *in vivo, in vitro* and *in silico* studies indicate pH gating of AQ(G)Ps and pin-point this effect to specific His residues as pH gates in the respective structures (55, 56, 89, 128, 139–152). Analyzing these publications, we were able to structurally separate five regions/positions in AQ(G)Ps where protonation/deprotonation of His residues was shown to modulate AQ(G)P function. We were overwhelmed by the fact that such a small and “simple” protein can be gated by pH via so many different ways, even though the molecular mechanisms of AQ(G)P modulation remains elusive in most of the referenced cases. One study on *Hs*AQP3 in lung cells indicated an additional His^2.-13^ (putative sixth gating site) in L1 at the periplasmic side (146), which we skipped in our analysis due to the fact that there is no high-resolution structure for AQP3 available, yet. **Figure 6** illustrates all 5 potential pH regulation sites in AQPs and AQGPs found in the literature (colored yellow and in green colors), mapped onto the *Ec*AQPZ structure as well as three additional His positions conserved in AQPs (shown in different blue tones). Interestingly, the latter histidines are found neither in any AQGP, nor in any AQPM. Only two pH gating sites have been structurally resolved so far, namely the pore exclusion of PIPs induced by His^5.-12^ in L5 (bright green, E) (56, 89) and a different side chain orientation in *Hs*AQP10 of the conserved His^h1.-3^ at the cytoplasmic end of the single-file pore (yellow, A) (55). The other positions in green color were located solely by computational and functional studies. Position A, His^h1.-3^, was located using MD simulations and experiments with *Hs*AQP5 (139) and *Hs*AQP4 (140, 141). Position D was identified in grapevine *Vv*TIP2;1 (151) using yeast cell assays and in *At*TIP5;1 (144) in oocytes. Position C is important in AQP0 (142, 143) as shown by pH dependent water measurements with oocytes. Position B was located in *Bt*AQP0 and *Rn*AQP4 using oocytes (143) in *Hs*AQP3 (146) in lung cells and in *Hs*AQP7 using yeast cells and MD simulations (150). **Figure S25** gives an overview of the number of His residues in the respective AQP structure and the occurrence in the respective potential pH gating site. A potential pH gating site which deserves detailed inspection is the pore lining His^h1.-3^ at the cytoplasmic entrance (within L2) labeled A and colored in yellow in **Figure 6A,** conserved in all investigated AQ(G)Ps except for *Kp*AQY1. MD simulations of *Hs*AQP5 (139) and *Hs*AQP4 (140, 141), experimental studies on *Hs*AQP4 (141) and *Hs*AQP10 (55) and structural investigations on *Hs*AQP10 (55) revealed this His residue as a pH gate. However, the function of several other AQPs carrying His in this position, including AQP1, were reported not to depend on pH. In an attempt to distinguish whether these differences result from inconclusive measurements or from differences in the local surroundings of the histidines, we have analyzed the H-bonds networks stabilizing His^h1.-3^ in the pore **(Figure 6B**). However, the amino acids involved in this network are largely conserved throughout the analyzed AQP structures (**Figure S26**). An H-bond network analysis similar to the one performed for the conserved Arg^h2.2^ in the selectivity filter revealed that His^h1.-3^ in position A is stabilized in 15 out of 20 structures at its side chain and in 19 out of 20 cases also at its backbone. However, we found no obvious correlation of the number of H-bonds stabilizing His^h1.-3^ itself or L2, housing His^h1.-3^, with AQ(G)Ps reported to be gated by pH at this position. Another explanation might be the variability in pKa values between 2.07 and 5.14 found using the program propka (153, 154) (**Figure S27**). Hence, shifted pKa values of the respective His residues may be responsible for the differences in pH gating. However, considering the limited accuracy of the pKa predictions and the limited number of functional studies, no obvious correlation could be described either.

**Figure 6.**
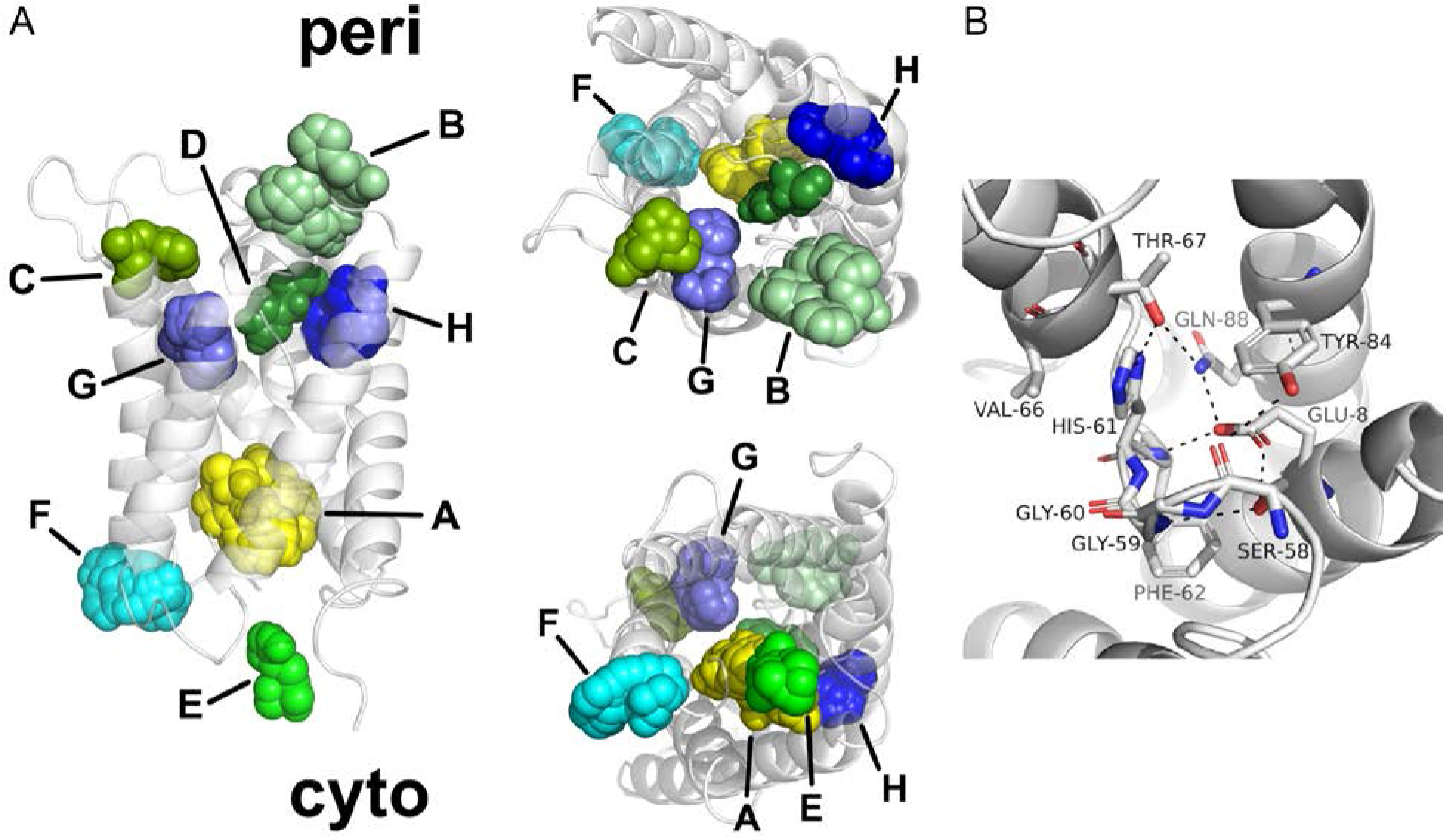
Possible pH gating sites and conserved His residues in AQ(G)Ps. **(A)** His positions among AQ(G)P structures potentially involved in pH gating shown in EcAQPZ. Color code according to the classification of individual positions listed in **Figure S25** and **Figure S27**. Please note, that in no AQ(G)P all here highlighted histidines are present at once. Occurrence of the respective His is 19, 5, 2, 1, 2, 8, 13 and 10 out of 20 for the positions A-H. Position A, His^h1.-3^, at the cytoplasmic side of the pore is conserved in all AQ(G)Ps except in KpAQY1. Position B, His^4.-14^ or His^4.-16^, in L4 at the periplasmic side appears in EcGLPF, OaAQP0, BtAQP0, HsAQP4, and RnAQP4. Positions C, His^2.-11^, in the periplasmic pore in L1, D, His^4-.18^, in L4 in the periplasmic pore and E, His^5.-12^, in L5 at the cytoplasmic mouth contain His only in AQP0s, AtTip2;1 and PIPs, respectively. Position F, His^h1.-8^, in the cytoplasmic vestibule is conserved in EcAQPZ, OaAQP0, BtAQP0, HsAQP1, BtAQP1, HsAQP2, HsAQP4 and RnAQP4. Position G, His^5.5^, is a part of the ar/R selectivity filter in all AQPs except AQGPs, AQPMs and AtTip2;1. Position H, His^6.-6^, in H6 at the periplasmic side is conserved in all classical AQPs and AtTip2;1 except for PIPs. In addition, periplasmic and cytoplasmic views are depicted. **(B)** Hydrogen bond network stabilizing the position of His61^h1.-3^ (position A, yellow) in L2 (EcAQPZ). Hydrogen bonds connecting Glu8^1.-6^ (2), Ser58^h1.-6^, Gly60^h1.-4^, Phe62^h1.-2^, His61^h1.-3^ (A), Thr67^h1.3^ (5), Tyr84^3.-7^ (6), and Gln88^3.-3^ (7) in sticks are indicated with dashed lines.

Interestingly, even though our structural analysis of the H-bond networks didn’t reveal any obvious indication why this His^h1.-3^ may gate water flux through some AQPs but not others (**Figure S24**), we found differences in His^h1.-3^ flexibility in our MD simulations. Whereas His^h1.-3^ formed stable H-bonds in *Bt*AQP1, *Ec*AQPZ and *Ec*GLPF, it was very mobile in *Hs*AQP4, hardly forming any H-bonds with neighboring residues (**Figure S23**). *Bt*AQP1, *Ec*AQPZ, and *Ec*GLPF showed significant H-bond probabilities with the conserved Glu^1.-6^ (position 2) in H1 and the conserved Thr^h1.3^ six positions towards the C-terminus located in HH1 (position 5 in Figure 9). In addition, all three AQ(G)Ps formed H-bonds with the conserved Ser^h1.-6^ in L2 three residues towards the N-terminus (position 2 in Figure 9), although only in the case of *Bt*AQP1 to a significant extend. Partly in line with these *in silico* results, our structural analysis revealed that the side chain of the conserved His^h1.-3^ forms either an H-bond with itself as in the case of *Bt*AQP1, the mentioned Thr^h1.3^ six positions towards the C-terminus located in HH1 (**Figure S26**) or its backbone with the conserved Glu^1.-6^ in H1. However, interactions with Ser^h1.-6^ in L2 are not obvious from the high-resolution structures. Intriguingly, a recent *in silico* study suggested the role of water coordination around His^h1.-3^ and interactions with the conserved Glu^1.-6^ in H1 in the gating behavior of *Hs*AQP10 (155). Similarly, we found major differences in the interaction with Glu^1.-6^, as outlined above and in the average number of H-bonds the His forms with pore water molecules (**Figure S20**). His95^h1.-3^ of *Hs*AQP4 forms at neutral pH on average 0.7±0.5 H-bonds with other pore lining amino acids, with 0.3±0.3 H-bonds to Glu^1.-6^ in H1 as calculated from the data presented in **Figure S23**. On the contrary, His76^h1.-3^ in *Bt*AQP1, His61^h1.-3^ in *Ec*AQPZ, and His66^h1.-3^ in *Ec*GLPF form 1.5±0.1, 2.0±0.1, and 1.9±0.04 H-bonds with other amino acids and 1.1±0.04, 1.1±0.03, and 1.0±0.02 H-bonds particularly with Glu^1.-6^, respectively. Vice versa, His95^h1.-3^ of *Hs*AQP4 formed on average 3.6±0.1 H-bonds with pore lining water molecules while His76^h1.-3^ in *Bt*AQP1, His61^h1.-3^ in *Ec*AQPZ, and His66^h1.-3^ in *Ec*GLPF formed only 2.3±0.1, 2.3±0.3, and 2.3±0.1, respectively (**Figure S20**). Hence, as our analysis of the crystal structures does not match the results from MD simulations, we suggest that His flexibility and pH gating reported in previous research may be due to differences in the vicinal water dynamics impinging on the stabilizing H-bonds of His^h1.-3^.

### 2.8. Roles and stability of the tetrameric fold of AQ(G)Ps

The AQ(G)P tetramers are stabilized via hydrophobic interactions at the protomer-protomer interface **(Figure 7A**). However, our analysis of all AQ(G)P interfaces revealed large differences in the shape and pattern of the hydrophobic protein-protein interfaces (**Figure S28**) compared to the rather uniform hydrophobic belt on the protein-lipid surface surrounding the tetrameric assembly (**Figure S29**). The central interfaces of the here studied AQ(G)Ps exhibit almost perfect surface complementarity with a differently pronounced dent at the center of the membrane towards the lipid bilayer. Additional stabilization is ensured by H-bonds and salt bridges. These are often necessary to imply specificity of the interaction interface (156). Salt-bridges at the protomer-protomer interface of the core proteins (i.e. proteins with truncated N- and C-terminus and without L1, L4, L5, and L7) are rare: only four AQ(G)Ps are stabilized by a salt-bridge in this region, and only *Af*AQPM and *Ec*AQPZ exhibit multiple salt-bridges, i.e. 3 and 2, respectively. Also, the full structures are stabilized by salt-bridges only in 8 out of 20 AQ(G)Ps. The number of hydrogen bonds at the protomer-protomer interface varies between 4 for *At*Tip2;1 to 15 for *Kp*AQY1. The situation changes when considering only the truncated protein versions (missing the N- and C-termini), with 13 remaining H-bonds for *Hs*AQP7 and *Hs*AQP10 and only 3 H-bonds stabilizing the tetramer for *At*Tip2;1 and *Kp*AQY1 (**Table S2**). While most of the formed H-bonds are also found within the truncated version of the AQ(G)Ps in Figure 9, some remarkable exceptions exist as illustrated by *Kp*AQY1 (out of 15 H-bonds only 3 are located at the truncated scaffold) and for *So*PIP2.1 (only 4 out of 10 H-bonds are found at the truncated proteins). The number of salt bridges varies between 0 (for most of the mammalian AQ(G)Ps) and 5 (for *Ec*AQPZ) in the full structures and 0 (for 16 structures) and 3 (for *Af*AQPM) for the truncated structures. Given those numbers, *At*Tip2;1 exhibits the lowest density of H-bonds per 1000 Å^2^ with only 2.3 and *Hs*AQP10 the highest with 9.5 for the full-length protein and 1.9 and 11 for the truncated protein respectively. The average AQ(G)P interface in our analysis has 5.2±1.6 H-bonds per 1000 Å^2^ and 0.4±0.7 salt bridges per 1000 Å^2^. The truncated AQ(G)Ps exhibit on average 5.0±2.2 H-bonds per 1000 Å^2^ and 0.2±0.5 salt bridges per 1000 Å^2^. Both values are at the lower end or even below published values for soluble protein interfaces of 5–10 hydrogen bonds per 1000 Å^2^ (157, 158) and around one salt bridge per 1000 Å^2^ (158). A detailed analysis of amino acids involved in the interaction at the protomer-protomer interface (**Figure S30**) revealed that H1, H2, the periplasmic end of H3, and the cytoplasmic end of H5 of one protomer interact with the periplasmic end of H2, H4, H5, and the cytoplasmic end of H6 of the neighboring protomer. In addition, the N- and C-terminus as well as L1, the N-terminal region of L4, L5 and the C-terminal side of L4 are involved in protomer contacts. While the interaction pattern is very diverse in the loops, it is more defined in the transmembrane region, with largely similar patterns but varying individual amino acids. This leads to an amino acid specific interaction strength via the number of hydrogen bonds, salt bridges, and the surface area involved in hydrophobic interactions (**Figure S30**).

**Figure 7.**
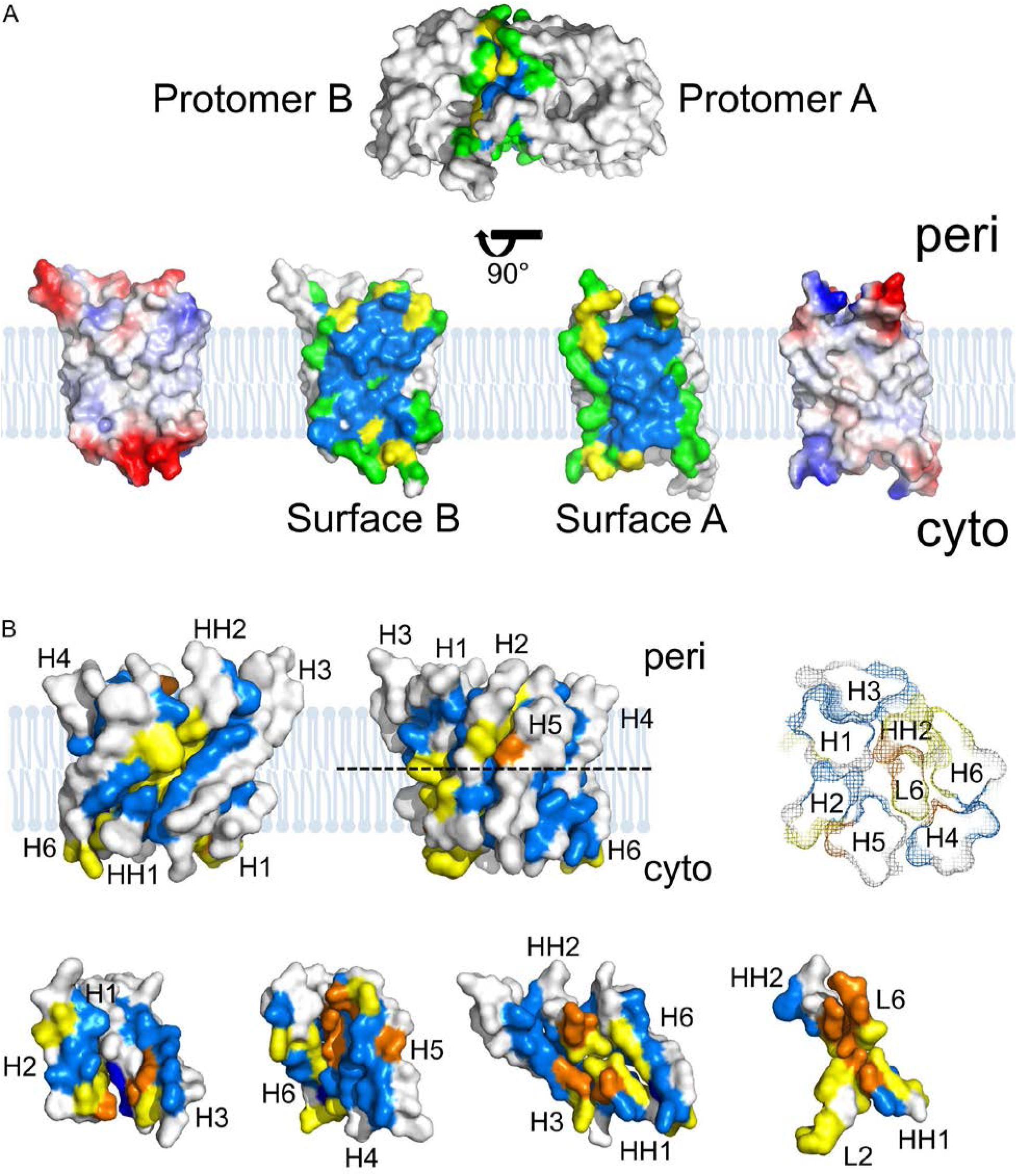
Exemplary inter- and intra-protomer interactions of EcAQPZ. **(A)** Hydrophobic residues that are involved in protomer-protomer surface interactions are colored in green (low degree of buried surface) and blue (high degree of buried surface). Residues involved in hydrogen bond formation are highlighted in yellow. The pictures at the very left and very right in the bottom row show net surface charge representations as estimated by PyMOL: white (no charge - hydrophobic), red (negative charge) and blue (positive charge) coloring. **(B)** Residues involved in hydrophobic interactions are colored in light blue (single interaction) or dark blue (multiple interactions). Residues forming hydrogen bonds are labeled in yellow (single hydrogen bond) or orange (multiple hydrogen bonds formed). The projection of a horizontal cut within the EcAQPZ protomer (dashed line) is shown in mesh representation at the top-right and the helix numbers are indicated (periplasmic view). Representative interaction surfaces among helices are shown in detail at the bottom.

Our analysis clearly shows that hydrogen bonds and salt bridges almost exclusively occur at protein-aqueous phase interface **(Figure 7A**, **Figure S28**). As there are hardly any methods available to get an idea about the relative contribution of each amino acid to the overall stability of the tetramer or the single protomers, we estimated buried surface areas (159) and interaction free energies ΔG of the respective interfaces using PDBePISA (160). Overall, ΔG is lower for mammalian AQPs as compared to archaea, bacteria, protozoa or yeast (**Table S2, Figure S31**). With an overall hydrophobic protein thickness of all AQ(Q)P structures of 30.2±1.0 nm as extracted from the OPM database (161) we didn’t find any dependence of ΔG on the potential membrane thickness (data not shown). Compared to the linear relation of protomer-protomer ΔGs on the buried surface area of the respective interface (**Figure S31**), there is no clear trend between inter- and intra-protomer ΔGs (**Figure S32**). Concomitant is no obvious trend for intra-protomer ΔGs on the AQ(G)P classes/subfamilies (**Figure S32**).

Here, we would like to note that depending on the method used to calculate these ΔG values, absolute values may vary. Consequently, using PDBePISA only allows us to rank relative differences between AQ(G)P structures. PDBePISA comprises any kind of contact interaction, including hydrogen bonds, into the ΔG calculation. However, as it assumes that potential hydrogen bonding partners become satisfied by hydrogen bonds to water upon dissociation/unfolding, this may significantly lower the contribution of hydrogen bonds on the overall interaction free energies of membrane proteins. As PDBePISA is designed for soluble protein complexes, neglecting the contribution and presence of the lipid bilayer during e.g. oligomer disruption into single protomers, this may lead to an underestimation of the contribution of H-bonds and salt bridges to the overall ΔG of membrane proteins. Experimental and theoretical studies suggest *E_hb_≈2–10* kcal/mol (162), whereas the remaining effect of decreased entropy of solvent due to the loss of mobility by bound molecules leads to an overall estimation of about *E_hb_≈0.6–1.5* kcal/mol per H-bond only (163, 164). Limited experimental data on the stabilization effect of salt bridges suggest that free energy contribution of a salt bridge is close to that of a hydrogen bond, amounting to *E_sb_≈0.9–1.25* kcal/mol (165, 166). A disulfide bond may contribute up to 2–8 kcal/mol (167, 168), yet, at lower occurrence. Hence, especially at the protomer-protomer interface of transmembrane proteins, hydrogen bonds might appear as major contributors into ΔG. This is a very interesting finding given the fact that AQPs exhibit on average less H-bonds per 1000 Å^2^ protomer-protomer interface than soluble proteins.

The residue specific analysis of intra-protomer helix-helix interactions reveals the key importance of a strong H-bond network anchoring L2 (connecting H2 and HH1), L6 (connecting H5 with HH2) and neighboring residues of the respective HH (**Figure S33**). These stabilizing H-bond networks, discussed in detail in the next chapter, ensure the desired selectivity and proper functionality of the protein, keeping the ar/R filter and NPA region in place. Moreover, L2 and L6 are inevitable in forming H-bonds by their backbone oxygens with pore water molecules, something not possible with a helical structure, where all backbone oxygens and amides are required to stabilize the secondary structure. Hence, structurally stable selectivity filters as well as pore lining (loop) residues are inevitable for proper function of AQ(G)Ps. As can be seen in **Figure 7B**, the rest of the helices surrounding the selectivity filter region are stabilized mainly by hydrophobic interactions (blue) with single H-bonds stabilizing the ends of the α-helices. Interestingly, AQGPs and AQPMs exhibit on average 4.3 and 4.0 stabilizing H-bonds between H1 and H3 and 2.5 and 2.0 H-bonds between H2 and H5 compared to 0.5 and 0.6 H-bonds in AQPs, respectively.

Thermostability assays with reconstituted *Rn*AQP4, *Ec*GLPF, *Mm*AQPM, and *Ec*AQPZ in *E. coli* total lipid extract revealed that these proteins turned inactive at 70 °C, 90 °C, 90 °C, and 100 °C, respectively (116). While *Ec*AQPZ revealed the steepest temperature dependence, *Ec*GLPF exhibited the flattest, which is reflected by the estimated thermal denaturation values (Tm) in Figure 8. A linear fit to the data of interaction free energies (**Table S2**) versus Tm values for these four AQ(G)Ps shows linear correlation of high credibility (Figure 8). This is surprising as PDBePISA ΔG values are thought to be a good relative approximation for hydrophobic interactions, underestimating the contribution of H-bonds or salt bridges in membrane proteins. Since neither the H-bonds or salt-bridges (8/0, 5/0, 8/1, and 6/5) nor the size of the protomer-protomer interaction interfaces (1488.6 Å^2^, 1545.3 Å^2^, 1763.4 Å^2^, and 1706.3 Å^2^) yield a similar correlation, we are led to the conclusion that hydrophobic interactions are the main determinant of thermal denaturation of AQ(G)Ps, with only a marginal contribution of H-bonds and salt-bridges. The later two are, as mentioned above, likely responsible for the specifity of the protomer-protomer interactions instead.

**Figure 8.**
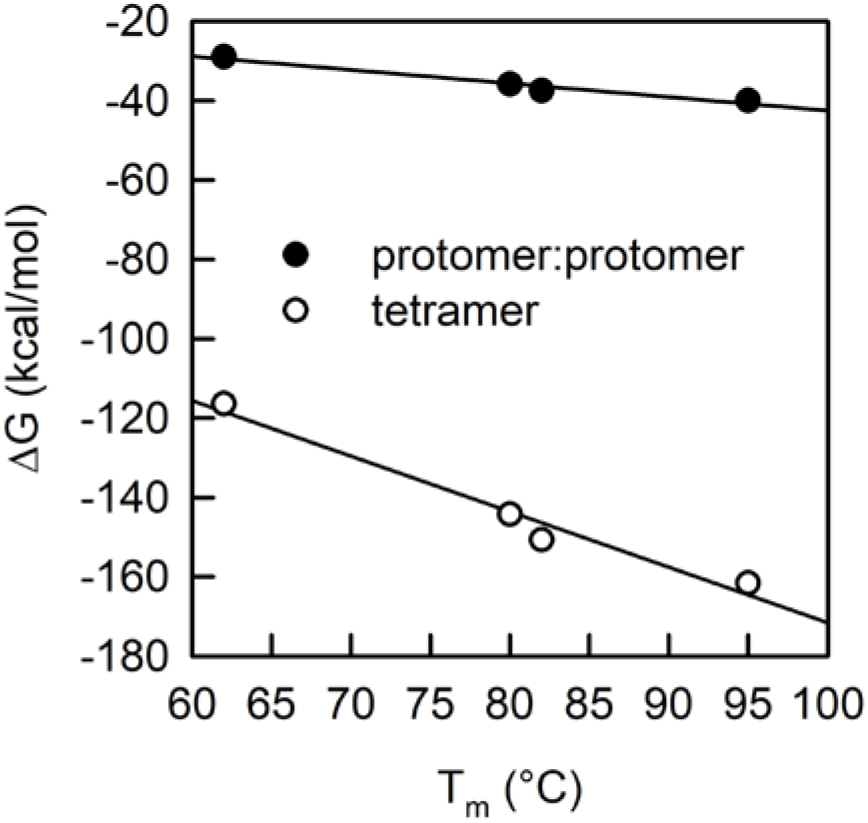
AQ(G)P thermostability. Thermal denaturation values (T_m_) in the order of RnAQP4 < EcGLPF < MmAQPM < EcAQPZ (116) are compared to the interaction free energies calculated by PDBePISA. The data is fitted with a linear regression resulting in k-values of −0.34 and −1.4 for the protomer-protomer (full circles) and tetramer (empty circles) case. Protomer-protomer ΔGs of −29 kcal/mol, −35.9 kcal/mol, −37.5 kcal/mol, and −40.0 kcal/mol and tetramer ΔGs of −116.4 kcal/mol, −144.3 kcal/mol, −150.7 kcal/mol, and −161.6 kcal/mol for RnAQP4, EcGLPF, MmAQPM, and EcAQPZ, respectively, were used according to **Table S2**.

### 2.9. Roles of highly conserved residues

In addition to the conserved Arg^h2.2^ in the ar/R selectivity filter including some of its interaction partners as well as the conserved His^h1.-3^ which we discussed in detail in section **2.7.1**. and **2.7.3**. and the well-known NPA motives, our sequence alignment (**Figure S1**) reveals additional highly conserved amino acids. For illustration purposes, we color coded these positions in our AQ(G)P scaffold (Figure 9). A closer look, shown in detail in **Figure S34**, discloses their diverse roles in stabilizing the AQ(G)Ps. The NPA motives (orange, **+**) connect the half helices HH1 and HH2 with the inner pore loops L2 and L6. The extended H-bond network connects the NPA motives to the transmembrane helical bundle of AQ(G)Ps. The inner pore loops L2 and L6 are stably anchored to the transmembrane helices by residues highlighted in magenta, in detail by Glu8^1.-6^ (**2**), Thr67^h1^.^3^ (**5),** Glu138^4.-4^ (**10**), Thr181^5^.^12^ (**13**), and Gln88^3.-3^ (**7**). Moreover, Gly59^h1.-5^ (**4**) shapes L2 via a loop internal interaction. Conserved Glys (labeled in dark blue) in the middle of helices ensure close H:H contact and include Gly91^3.0^ (**8**), Gly167^5.-2^ (**12**), and Gly215^6.3^ (**16**). These Glys are parts of **G**(A/S)XXXG(A), P(G/S/A)XXX**G** and G(A/V)XXX**G**XXXG(A/F) motives similar to the well-known glycine zippers (169). Lipid interactions or larger spacer residues are labeled in cyan or grey (conserved in AQGPs, only) and include lipid-binding Arg3^1.-11^ (**1**), spacers Tyr84^3.-7^ (**6**), Phe/Tyr208^6.-4^ (in case of AQGPs Pro^6.-4^) (**20**) and Tyr223^6.11^ (**17**), as well as membrane anchors (170) Trp206^6.-6^ (**14**), Phe98^3.2^ (**18**) and Phe149^4.-7^ (**19**). Moreover, Pro212^6.0^ (**15**) and neighboring hydrophobic AAs are part of the double glycine zipper motive in H6 enabling close interaction with H4 and HH2. Other conserved positions are labeled in green: Gly21^1.7^ (**3**) responsible for a kink in H1 which is, however, only obvious in 7 structures (*Ec*GLPF, *Ec*AQPZ, *Af*AQPM, *Mm*AQPM, *At*AQPZ, *At*TIP2;1, *Bt*AQP1). Ser130^4.-12^ (**9**) at the crossover between L4 and H4 stabilizes the oligomeric assembly via a backbone interaction to H3 of the neighboring protomer in AQPZ, and Ser142^4.0^ (**11**) (Thr^4.0^ in all structures except of *Ec*AQPZ) in H4 stabilizing the connection with H6, keeping the cytoplasmic side of these helices in place.

**Figure 9.**
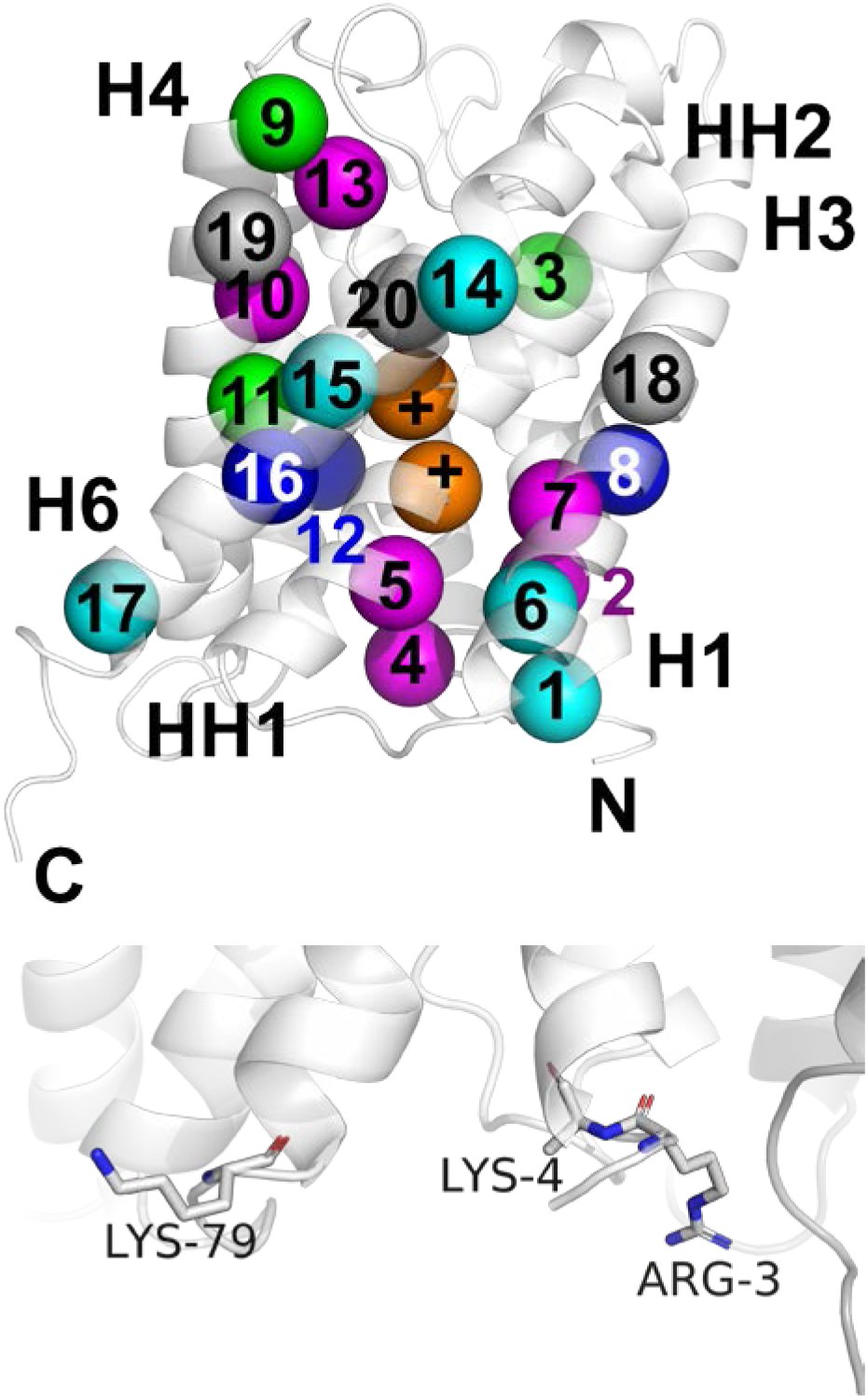
Location of conserved amino acid residues in the AQ(G)P scaffold. 21 locations are delineated in the sequence alignment in **Figure S1**: Arg3^1.-11^ (**1**, 65% conserved & 95% charge similarity), Glu8^1.-6^ (**2**, 100% conserved), Gly21^1.7^ (**3**, 85% conserved), Gly56^h1.-5^ (**4**, 100% conserved), Thr67^h1.3^ (**5**, 85% conserved), Tyr84^3.-7^ (**6**, 90% conserved), Gln88^3.-3^ (**7**, 100% conserved), Gly91^3.0^ (**8**, 90% conserved), Ser130^4.-12^ (**9**, 40% conserved & 90% type similarity), Glu138^4.-4^ (**10**, 95% conserved), Thr142^4.0^ (**11**, 95% conserved), Gly167^5.-2^ (**12**, 100% conserved), Thr181^5.12^ (**13**, 85% conserved), Trp206^6.-6^ (**14**, 85% conserved), Pro212^6.0^ (**15**, 100% conserved), Gly216^6.3^ (**16**, 100% conserved), Tyr223^6.11^ (**17**, 80% conserved & 100% steric similarity), Asn63^h1.-1^ and Asn186^h2.-1^ as part of the NPA motives (**+**, 100% conserved), Phe98^3.2^ (**18**, 100% conserved in AQGPs) and Phe149^4.-7^ (**19**, 100% conserved in AQGPs), Pro236^6.-4^ (**20**, 100% conserved in AQGPs). Snapshots of the H-bonding networks of these conserved residues are depicted in **Figure S34**. Conservation is calculated as probability occurrence of the very same AA in all 20 high-resolution structures. Lipid interactions or larger spacer residues are labeled in cyan, the NPA motive in orange, residues fixing HHs and loops to HHs in pink, Glys and Alas for close H:H contact in dark blue, and other special positions in green. **Bottom:** Arg/Lys (**1**) at the groove between two protomers (light and dark grey) represent a conserved lipid interaction site. Shown are the Arg3^1.-11^ and Lys4^1.-10^ at the cytoplasmic end of H1 and Lys79^3.-12^ at the cytoplasmic end of H3 of EcAQPZ.

Taken together, AQ(G)Ps contain multiple conserved residues whose art, localization, and H-bonding networks hint to very diverse roles, some of which are open for experimental validation.

#### 2.9.1. Lipid interaction sites with negatively charged lipids are conserved

*Ec*AQPZ is significantly stabilized by cardiolipin (CL) as shown by mass spectrometry (171). In addition, it was discovered that CL binds at the contact sites of protomers in the tetrameric assembly (172). Investigating CL interactions utilizing a multitude of *E. coli* inner membrane proteins *in silico* identified a typical CL binding site to harbor two to three basic residues in close proximity, at least one polar residue, and one or more aromatic residues slightly deeper within the membrane (173). Similar to *Ec*AQPZ, CL was also found to bind preferentially to this cytoplasmic crevice at the contact of two neighboring *Bt*AQP1 protomers, as evident from MD simulations using a native-like model of *E. coli* polar lipid extract (174). In the latter study, the rest of the cytoplasmic protein surface was covered with negatively charged phosphatidylglycerols, which were attracted by a number of positively charged residues (Lys7^1.-16^, Lys8^1.-15^, Arg12^1.-11^, Arg95^3.-11^, Arg243^6.25^, and Lys245^6.27^), similarly to CL. Our analysis here reveals that the above-mentioned position of positively charged residues at the contact site between AQP protomers is conserved. **Figure 9** illustrates the relative orientation of positive residues Arg3^1.-11^ and Lys4^1.-10^ at the cytoplasmic end of H1 and Lys79^3.-12^ at the cytoplasmic end of H3 of *Ec*AQPZ. Both locations are equipped with positive charges in all AQPs under investigation (**Figure S1**). Positively charged residues have been shown before to specifically target proteins to negatively charged surfaces of diverse membranes (175) and to control the orientation of the protein upon insertion into the lipid membrane (176). The above mentioned conserved positively charged residues at the mouth of the crevice between two AQ(G)P protomers hint to the fact that AQ(G)Ps might additionally use the high abundance of negatively charged lipids at the cytoplasmic side of the membranes to stabilize their tetrameric structure. Yet, the absolute impact of specific lipid interactions of an AQ(G)P embedded in a native lipid bilayer on AQ(G)P stability has not yet been addressed.

### 2.10. The central pore

The biological significance of AQ(G)P tetramerization is still elusive. In general, oligomerization might lead to structural and/or proteolytic stability, functional diversity, regulatory mechanisms, and formation of binding cavities (177). One side-effect of AQ(G)P tetramerization is the formation of a potential central pore. Several studies suggest ion (178–181) and gas (182) transport through the central, potential pore. As the topic is highly debated (75, 110, 111), yet biological significance is still missing, we have conducted an analysis of the geometrical and chemical properties of these potentially channel forming central pores. In comparison to the protomeric pores, most of the profiles of the central channel reveal constrictions of ≤ 0.65 Å radius, which is geometrically too narrow for water molecules to pass (**Figure S35**) (183). Notable exceptions are *Bt*AQP0, *Bt*AQP1, *Hs*AQP1, and *Hs*AQP2 with 0.7 Å, *Mm*AQPM and *At*Tip2;1 with 0.9 Å, *Hs*AQP4 and *Rn*AQP4 with 0.9-1 Å, and *Ec*GlpF with 1 Å. **Figure S36** illustrates the plethora of different pore geometries found for our structural dataset. Compared to the protomeric pore geometries, the shapes are much more diverse and less conserved. The considerable differences in pore length are partly caused by significant differences in the length of the cyto-/periplasmic loops connecting the transmembrane helices. *Pf*AQP is completely closed at the cytoplasmic side.

However, the hydrophobic characteristic of the inner surface of the central pores is conserved among all analyzed AQ(G)Ps: Except for a ring of charged residues at the periplasmic side of the channel in case of *Ec*GLPF, *Pf*AQP and mammalian AQP1s, the narrow pore is solely decorated by hydrophobic side chains (**Figure S37**), which renders potential water passage through the central pore illusive. This presumption could be confirmed by analyzing water permeation events using MD simulations, where 0.99% (11 versus 1106), 0.96% (16 versus 1663), 2.49% (9 versus 361), and 0.43% (6 versus 1135) of the water molecules passed the central pore as compared to the 4 protomer pores within a simulation time of 500 ns for *Bt*AQP1, *Ec*AQPZ, *Hs*AQP4, and *Ec*GLPF, respectively. Interestingly, narrow (2 Å diameter) carbon nanotubes (CNTs) are thought to be highly water permeable despite their hydrophobic nature (184). This discrepancy may partly be due to different pore surfaces, which are homogeneous in CNTs and highly inhomogeneous in AQPs. In addition, CNTs are decorated with benzenes compared to methyl groups in AQPs. Moreover, a ring of negatively charged residues may serve as a putative divalent cation binding spot as found for Glu43^2.-8^ in *Ec*GLPF (185), further obstructing the pore for other solutes (**Figure S37**). An alanine mutant (Glu43A1a^2.-8^) suggested the involvement of the respective Mg^2+^ binding site in the overall stability of *Ec*GLPF, shifting the oligomeric state dramatically towards the monomer (186). Stabilization of the tetramer in the presence of Mg^2+^ ions was also reported by Borgnia at al. (187). However, the effect could not be confirmed in experiments with EDTA as a chelator (188). In addition to Glu43^2.-8^, the crystal structure of *Ec*GLPF localized a second Mg^2+^ next to Trp42^2.-9^ (43). Glu43^2.-8^ might also be involved in the binding of specific lipids *in vivo* (43, 53). In wider central channels, i.e. the hexameric *Hp*UreI, pore obstruction by lipids is a common phenomenon (96). In the yeast aquaporin, *Kp*AQY1, another finding suggests chloride ions to be bound in the central channel next to Trp^5.-9^ (*115*). Generally, structural comparison with narrow ion channels reveals that their ion selective pores are not hydrophobic but decorated with countercharges in respect to the transported ion permeating the channel (189, 190). Thus, the hydrophobic central pore of AQ(G)Ps is unlikely to efficiently transport water or small ions even if the narrow constriction would theoretically provide enough space.

## 3. Conclusions and Outlook

With the overall structure of aquaporins being well established, including a sub-Ångström structure of *Kp*AQY1 *(44)*, recent structures combined with experiments and simulations (*54, 55, 57*) have provided great insight into the transport, selectivity, and regulation mechanisms of individual AQPs. Here, we took a new approach, analyzing all 20 non-redundant high-resolution structures deposited in the protein database to unravel structural peculiarities of AQ(G)Ps hidden by the analysis of single AQP structures or limited subsets.

In principle, AQ(G)Ps constitute simple proteins, with an equal fold of 6 transmembrane helices and 2 half-helices, nevertheless capable of fulfilling diverse function, regulation, and selectivity. First, we want to elaborate on the question how it is possible that seemingly highly similar proteins exhibit such a wide variance in unitary permeability values (*p_f_*), as shown in Figure 1. The major determinants of single-file transport of water through narrow membrane channels were suggested to be the number of H-bonds water molecules may form with pore lining residues (4), channel gating by pore lining residues (77), positive charges at the pore mouth potentially reducing the dehydration penalty (81) and possibly the shape of the entry/exit vestibules (82). Moreover, the geometry of the pore as well as structural changes due to lipid interactions may depict additional determinants. The characteristic hourglass shape of AQPs was found to be the optimum for a hydrodynamic dissipation process, maximizing channel permeability (82). However, as it can be seen from **Figure 3, S8** and **S9**, the shape and angle of the hourglass shaped pore entrances are very conserved among AQ(G)Ps, thus leaving hardly any room to explain permeability differences among AQ(G)Ps. Similarly, the overall pore geometry (width) could potentially explain *p_f_* differences between AQPs and AQGPs but hardly among AQPs, as crucial differences in pore geometry within this group are missing (**Figure S10**). The effect of positive charges at the pore entrance potentially reducing the dehydration penalty is thought to be minor (81). The latter estimate expanded to the current set of 20 AQ(G)P structures (**Figure S8**) reveals comparable differences in the number and distribution of positive AAs next to the pore entrances. Similarly, our structural analysis suggests that the potential number of H-bonds water molecules may form with pore lining residues is hardly variable. H-bonds formed to single-file water molecules are either formed via side chains of conserved residues in the NPA motives or in the ar/R filter or via backbone oxygens of L2 and L6. Interestingly, our MD simulations depict significant variations in the overall number of H-bonds single-file water molecules form with these conserved pore lining residues among different AQ(G)Ps, which would principally enable a certain variability in p_f_. What is left are gating effects by pore lining residues (e.g. the phenolic barriers drastically reducing the permeability of AQP0 (47)) or lipid specific effects.

In line with these insights, our structural analysis and MD simulations clearly show that AQ(G)Ps do not exhibit a universally open pore. In contrast, water flow through the single-file pores is modulated by pore lining residues as sporadically already mentioned in literature. Even though the physiological significance of water flux modulation via the conserved His at the cytoplasmic side of the single-file region can be envisioned similar to pH regulated human adipose glycerol flux through *Hs*AQP10 (55), the picture is less obvious regarding gating of the conserved Arg^h2.2^ of the ar/R selectivity filter. Theoretically the position of the Arg^h2.2^ sidechain in the pore could be regulated via the transmembrane potential, lipid asymmetry, or binding of highly charged soluble proteins to the pore entrance, the latter two impinging on the membrane potential. So far, such an effect on water permeability through *Hs*AQP1 and *Hs*AQP4 was only seen *in silico* at unphysiologically high membrane potentials of >±0.5V (134). However, high transmembrane potential can lead to electroporation of the membrane (191, 192) or to denaturation of transmembrane proteins (193). The importance of side chain fluctuations of Met^h2.-2^ positioned in the direct vicinity of the NPA and ar/R motives awaits clarification. Are they a result of the imperfection of the respective AQP or part of an elaborate regulation mechanism? This is a question which is not easy to answer. However, what we could show for the dataset analysed herein by comparing the H-bond network stabilizing the Arg^h2.2^, it is possible to predict its mobility *in-silico* with a clear impact on water passage through the pores (**Figure S21-S24**).

Which structural effects specific lipid interactions have on AQ(G)P functionality and how such direct structural regulatory mechanisms would look like, remains an open challenge. Nonetheless, as it is known that specific binding of lipids stabilizes the oligomeric assembly (159, 194), we speculate that stabilization of the oligomeric assembly has an impact on the flexibility of pore lining residues. This could change the probability of pore lining residues to reside within the channel pore obstructing it for a certain percentage of time as well as potentially influence its selectivity. Hence, whereas the maximum water permeability of AQ(G)Ps is defined by the pore characteristics discussed above, we speculate that the net flux through an AQ(G)P is defined by channel gating via flexible pore lining residues. Examples could be a strongly modulated ribitol transport capability of *Ec*GLPF by negatively charged lipids (195) and an increased *Ec*AQPZ water permeability after CL binding (171).

Substrate discrimination in water channels is thought to depend on a complex interplay between the solute, pore size and polarity, with the pore size determining the exclusion properties but not solute selectivity (64). In accordance, the pore size of AQGPs is larger throughout the pore compared to AQPs and AQPMs (Figure 3, **Figure S10**). However, so far, it was not clear if this larger pore size is solely constituted by pore lining residues or if the AQGP scaffold is distinctly different from that of AQPs. **Figure S15** and **Figure S17** visualize that this is indeed the case. AQGPs do not only exhibit a larger protomer-protomer distance by 1.1 Å and an overall increased dimension of the tetrameric assembly by 2.6 Å along the protomer-lipid interface, but also the relative distances of helices in the protomer are increased. Thereby, H5 localization further away from the protomer pore goes hand in hand with an increased H4:H5 distance. The latter is possible due to an evolutionary groove at the H4:H5 interface, with perfectly parallel oriented helices within the protomeric arrangement (**Figure S38**). In contrast, all other helix-helix interactions are much more refined, with the helices tilted to each other forming helix-helix cross overs with highly conserved Gly and Ala residues enabling close helix-helix contacts. Concerning AQPMs, we showed that despite having a broader pore than AQPs, yet with a rather similar scaffold size, it is possible to reach glycerol selectivity with an AQP scaffold but at the expense of efficient permeability. To achieve both, a special set of pore lining residues and an adequately expanded scaffold as in AQGPs seem mandatory.

Our analysis reveals internal H-bond network stabilizing both HH and adjacent loops L2 and L6, which harbor most of the H-bonds donating and accepting residues in the single-file region of AQ(G)Ps. Also, the H-bond network shapes the structure and the dynamics of the ar/R filter and thus its selectivity **(Figure 7B**, **Figure S33**). Together with the strong conservation of the involved residues (Figure 1, **Figure S34**) this highlights the importance of a stable single-file region for AQ(G)P selectivity and permeability. We speculate that the tetrameric arrangement is inevitable to ensure this structural necessity as monomeric *Ec*AQPZ (196) and *Ec*GLPF (197–199) exhibited reduced activities in terms of water and ribitol flux, respectively. Monomeric *Ec*GLPF was also less resistant to proteolysis in *E. coli* (186), with a significantly stabilized tetrameric assembly *in vivo* (188). Additional stabilization via specific lipid interactions within the complex natural lipid composition (174) may have evolved in parallel to inter- and intra-protomer evolution. In any case, optimization of tetramer stability is a tradeoff with protein aggregation and protein folding, as protomer stability in the lipid bilayer is a prerequisite for AQ(G)P folding into the membrane and consequent tetramerization. This protomer pre-insertion into the membrane may explain why AQ(G)P protomers exhibit hardly any polar residues at the protomer-protomer interface within the membrane (**Figure S28**). Instead, the oligomeric assembly is mainly stabilized via hydrophobic interactions within this region, supported by lipid interactions, H-bonds, and salt-bridges. The latter located mostly at the lipid bilayer to aqueous phase interfacial regions. Even though the individual contributions to tetramer stability still stay elusive, the here discovered correlation of the thermal denaturation temperatures for four AQ(G)Ps with the interaction free energies calculated by PDBePISA imply the crucial role of hydrophobic interactions for the stability of the AQ(G)P fold.

AQ(G)Ps are seen as potential building blocks of next generation filter membranes (17–28). Furthermore, they are treated as the “holy grail” of native water conducting pores, due to their superior performance in respect to their high permeability and great selectivity. Hence AQ(G)Ps serve as templates for a whole community designing artificial water channels (30–37). Desalination and distillation are among the most resource intensive industrial processes (200). Therefore, an enormous effort is spent to develop filter membranes with increased energy efficiency. As a first step, functional *Ec*AQPZ was already successfully incorporated into polymer vesicles and membranes (201), with already some successful commercialization of Aquaporin Inside™ desalination membranes from Aquaporin A/S (202). Still, this idea is afflicted with the prejudice that AQ(G)Ps are not stable enough (17) and might degrade in the presence of harsh conditions used in course of industrial membrane formation. This may be even more relevant when using other AQ(G)Ps, which are inherently less stable than *Ec*AQPZ, to expand the range of applications from pure desalination or water filtration to biotechnological applications like small molecule recovery. Our rigorous structural analysis is a first step towards the goal to create bio-inspired AQ(G)P variants with optimized stability and tuned selectivity for next generation biomimetic separation membranes. An ideal water channel in this respect shall combine high permeability, perfect selectivity, and exceptional structural stability, as to withstand adverse physical and chemical conditions. Altering the amino acid sequence of AQ(G)Ps in order to optimize the key aspects mentioned above, requires comprehensive understanding of the underlying structural and functional features. Consequently, this work can be used to choose the most appropriate scaffold in terms of AQ(G)P function and stability and combine interaction hotspots, purely hydrophobic in nature or specific via H-bonds or salt-bridges, found in other AQ(G)Ps into the chosen scaffold.

## 4. Material and Methods

### 4.1. Analysis and preparations of the AQ(G)P structures

The PDB database was surveyed for all currently available AQP and AQGP structures. The resulting comprehensive list, stating the respective resolution, method of structure elucidation as well as year of submission and reference to the original publication, if available, can be found in the appendix (**Table S1**).

For the analysis presented in this paper, we generated a non-redundant selection of AQ(G)P structures, by exclusion of identical, marginally mutated, or substrate containing structures of the same organism. The targets were chosen based on their apparent resolution (the higher the better) and the absence of mutations (wild type structures were preferred). The resulting list contains 20 structures: 2 AQPM, a subclass of archaeal AQPs first discovered in *Methanothermobacter marburgis*, 3 bacterial AQ(G)Ps, 2 structures originating from protozoa or yeast, 3 plant aquaporins, and 10 mammalian AQ(G)P structures. The corresponding amino acid sequences were aligned in Jalview (version 2.10.5)(203) utilizing the ClustalOmega algorithm with default settings and subsequently used for the visualization of shared AQ(G)P features.

To obtain root mean square deviation (RMSD) values for structure alignments of wildtype AQ(G)Ps, we used the standard alignment function of PyMOL (Schrödinger, version 2.3.2). *Ec*AQPZ (PDB: 1RC2) served as a reference for most of the following analysis due to its high resolution and most compact structure, with minimal loops, and N- and C-termini. Furthermore, it shows an astonishing stability(116) rendering *Ec*AQPZ promising candidate for biotechnological applications (201, 204). N- and C-termini of all aquaporins were removed according to the remaining quality of the alignment. The resulting truncated models were named the AQ(G)P “core”. Helix and loop lengths, as well as the respective amino acid composition of the chains were analyzed in PyMOL. In the core models loops 1, 4, 5 and 7 were omitted due to their high flexibility/variability. The C_α_ RMSD values of the core structures relative to *Ec*AQPZ (1RC2) were calculated using the PyMOL align command, with outlier rejection turned off. Next to RMSD estimation for each C_α_ also RMSDs of C_α_ in five distinct horizontal plains along the z-axis of the AQ(G)Ps were estimated.

The PDBePISA server (160, 205) was used to calculate interaction areas and the gain in solvation energy upon interface formation, ΔGs, between AQ(G)P protomers as well as internally, between individual helices and loops. Furthermore, detailed analysis of individual interactions derived from PDBePISA provided information about amino acids involved in hydrophobic interactions (buried area percentage of interfacing residues), hydrogen bonds, and salt bridges. The detailed interaction profiles were registered to a template sequence alignment of the 20 AQ(G)P structures and thus gave rise to detailed depictions of the inter- and intra-molecular interactions within these structures.

The shape of AQ(G)P protomer channels as well as the shape of the central pore in AQ(G)P tetramers were analyzed with the program HOLE(123) utilizing the xplor algorithm and providing coordinates of a single water molecule in the center of the aligned AQ(G)P structures to serve as a start point for the channel traces.

The pKa values of histidine residues were evaluated using the program propka (version 3.4.0)(153, 154).

All structure figures were prepared in PyMOL(206). Calculations were performed in Excel and graphs plotted using the program VEUSZ (Jeremy Sanders, version 2.1.1) and R (207).

Evolutionary analysis was conducted in MEGA11 (208). The phylogenetic tree with the highest log likelihood (−4884.76) is shown in **Figure S3**. Initial tree(s) for the heuristic search were obtained automatically by applying Neighbor-Join and BioNJ algorithms to a matrix of pairwise distances estimated using the JTT model, and then selecting the topology with superior log likelihood value. This analysis involved 20 amino acid sequences. There were a total of 168 positions in the final dataset.

### 4.2. Molecular dynamics simulations of AQ(G)Ps

Molecular dynamics (MD) simulations were performed using the GROMACS package version 2016.3 and 2018.6(209). In each simulation system a tetrameric AQ(G)P was embedded in an *E. coli* polar lipid extract membrane model “Avanti”(210) containing 14 different lipid types by our sequential multiscaling methodology(211) as described in (210). The membrane consisting of 384 lipids, was solvated by ~29 000 water molecules and counter Na^+^ ions. The tetrameric *Bt*AQP1, *Hs*AQP4, *Ec*GLPF, and *Ec*AQPZ were modelled based on the crystal structures 1J4N (49), 3GD8 (51), 1FX8 (43), and 2ABM (136), respectively. Each chain of *Bt*AQP1 contained residues M1-S249, the shortened C-terminus was thereby capped by an amine group. Each *Hs*AQP4 chain consisted of residues Q32-P254 and each *Ec*GLPF chain was built of T6-E267. In both, *Hs*AQP4 and *Ec*GLPF the shortened N-termini carried an NH2 group and the shortened C-termini were capped by an amine group. Each *Ec*AQPZ chain contained all *Ec*AQPZ residues, i.e. M1-D231 and thus both termini were charged. The histidine in the ar/R filter of all three AQPs was protonated on N^δ^ (212), due to its higher water permeability. All other histidines were protonated on N^ε^. All titratable amino acids were protonated according to their preferred protonation state at pH 7. The 500 ns long production molecular dynamics simulations were performed at 296 K, using the CHARMM36m force field(213–215) and TIP4p water model(216). The trajectories were written out every 2 ps. For more details see (210).

The estimation of the number of water molecules permeating the AQ(G)P pores was performed by g_flux(217) through a cylinder with a radius of 0.75 nm and a length of 2 nm. Analysis of hydrogen bonds, side chain angles and distances between residues succeeded by standard GROMACS analysis tools.

## Supplementary Information

All Tables are shared as excel files.

## Acknowledgements

The financial support for this study is from the Austrian Science Fund (P 35541) to AH. The molecular dynamics simulations in this work were performed on the supercomputers ForHLR and BwForCluster BiNAC funded by the Ministry of Science, Research and the Arts Baden-Württemberg and by the Federal Ministry of Education and Research. K.P. was funded by Deutsche Forschungsgemeinschaft (DFG, German Research Foundation) under Germany’s Excellence Strategy - EXC 2075 – 390740016. KP also acknowledges the support by the Stuttgart Center for Simulation Science (SimTech).

## Author contributions

N. G-M., C. S., L. U., S. K. N. T., and A. H. analyzed AQ(G)P structures. K. P. performed and analyzed the molecular dynamics simulations. N. G-M., C. S., K. P and A. H. conceptualized the study. N. G-M., C. S., K. P and A. H. interpreted the results. N. G-M., C. S., K. P and A. H. wrote the manuscript. All authors approved the manuscript.

## References

1. Preston GM, Carroll TP, Guggino WB, & Agre P (1992) Appearance of water channels in Xenopus oocytes expressing red cell CHIP28 protein. Science 256(5055):385–387.

2. Zeidel ML, Ambudkar SV, Smith BL, & Agre P (1992) Reconstituion of functional water channels in liposomes containing purified red cell chip28 protein. Biochemistry 31:7436–7440.

3. Kreida S & Toernroth-Horsefield S (2015) Structural insights into aquaporin selectivity and regulation. Curr. Opin. Struct. Biol. 33:126–134.

4. Horner A, et al. (2015) The mobility of single-file water molecules is governed by the number of H-bonds they may form with channel-lining residues. Sci Adv 1(2):e1400083.

5. Fu D, Libson A, & Stroud R (2002) The structure of GlpF, a glycerol conducting channel. Novartis Found Symp 245:51–61; discussion 61-55, 165-168.

6. King LS, Kozono D, & Agre P (2004) From structure to disease: the evolving tale of aquaporin biology. Nature Reviews Molecular Cell Biology 5:687.

7. Abascal F, Irisarri I, & Zardoya R (2014) Diversity and evolution of membrane intrinsic proteins. Biochimica et Biophysica Acta (BBA) - General Subjects 1840(5):1468–1481.

8. Deshmukh RK, et al. (2015) A precise spacing between the NPA domains of aquaporins is essential for silicon permeability in plants. The Plant Journal 83(3):489–500.

9. Verkman AS (2012) Aquaporins in clinical medicine. Annu. Rev. Med. 63:303–316.

10. Papadopoulos MC & Saadoun S (2015) Key roles of aquaporins in tumor biology. Biochim. Biophys. Acta 1848(10 Pt B):2576–2583.

11. Azad AK, et al. (2021) Human Aquaporins: Functional Diversity and Potential Roles in Infectious and Non-infectious Diseases. Front Genet 12:654865.

12. Verkman AS, Anderson MO, & Papadopoulos MC (2014) Aquaporins: important but elusive drug targets. Nat. Rev. Drug Discov. 13(4):259–277.

13. Papadopoulos MC & Verkman AS (2013) Aquaporin water channels in the nervous system. Nature Reviews Neuroscience 14:265.

14. Maurel C, Verdoucq L, Luu DT, & Santoni V (2008) Plant aquaporins: membrane channels with multiple integrated functions. Annu. Rev. Plant Biol. 59:595–624.

15. Shekoofa A & Sinclair T (2018) Aquaporin Activity to Improve Crop Drought Tolerance. Cells 7(9):123.

16. Zargar SM, et al. (2017) Aquaporins as potential drought tolerance inducing proteins: Towards instigating stress tolerance. Journal of Proteomics 169:233–238.

17. To J & Torres J (2015) Can stabilization and inhibition of aquaporins contribute to future development of biomimetic membranes? Membranes 5(3):352–368.

18. Song W, Lang C, Shen Y-x, & Kumar M (2018) Design Considerations for Artificial Water Channel–Based Membranes. Annual Review of Materials Research 48(1):57–82.

19. Ren T, et al. (2017) Membrane Protein Insertion into and Compatibility with Biomimetic Membranes. Adv Biosyst 1(7):e1700053.

20. Abdelrasoul A, Doan H, Lohi A, & Cheng C-H (2018) Aquaporin-Based Biomimetic and Bioinspired Membranes for New Frontiers in Sustainable Water Treatment Technology: Approaches and Challenges. Polymer Science, Series A 60(4):429–450.

21. Fuwad A, Ryu H, Malmstadt N, Kim SM, & Jeon T-J (2019) Biomimetic membranes as potential tools for water purification: Preceding and future avenues. Desalination 458:97–115.

22. Li XS, et al. (2012) Preparation of supported lipid membranes for aquaporin Z incorporation. Colloids and Surfaces B-Biointerfaces 94:333–340.

23. Tang C, Wang Z, Petrinić I, Fane AG, & Hélix-Nielsen C (2015) Biomimetic aquaporin membranes coming of age. Desalination 368:89–105.

24. Qi S, et al. (2016) Aquaporin-based biomimetic reverse osmosis membranes: Stability and long term performance. J. Membr. Sci. 508:94–103.

25. Li Y, Qi S, Tian M, Widjajanti W, & Wang R (2019) Fabrication of aquaporin-based biomimetic membrane for seawater desalination. Desalination 467:103–112.

26. Wagh P & Escobar IC (2019) Biomimetic and bioinspired membranes for water purification: A critical review and future directions. Environmental Progress & Sustainable Energy 38(3).

27. Liang Z, et al. (2019) Performance evaluation of interfacial polymerisation-fabricated aquaporin-based biomimetic membranes in forward osmosis. RSC Advances 9(19):10715–10726.

28. Martinez-Ballesta MC, Garcia-Ibanez P, Yepes-Molina L, Rios JJ, & Carvajal M (2018) The Expanding Role of Vesicles Containing Aquaporins. Cells 7(10).

29. Song W & Kumar M (2019) Artificial water channels: toward and beyond desalination. Current Opinion in Chemical Engineering 25:9–17.

30. Shen YX, et al. (2018) Achieving high permeability and enhanced selectivity for Angstrom-scale separations using artificial water channel membranes. Nat Commun 9(1):2294.

31. Song W, Tu YM, Oh H, Samineni L, & Kumar M (2019) Hierarchical Optimization of High-Performance Biomimetic and Bioinspired Membranes. Langmuir 35(3):589–607.

32. Shen YX, et al. (2015) Highly permeable artificial water channels that can self-assemble into two-dimensional arrays. Proc Natl Acad Sci U S A 112(32):9810–9815.

33. Kocsis I, Sun Z, Legrand YM, & Barboiu M (2018) Artificial water channels—deconvolution of natural Aquaporins through synthetic design. npj Clean Water 1(1):13.

34. Murail S, et al. (2018) Water permeation across artificial I-quartet membrane channels: from structure to disorder. Faraday Discuss. 209(0):125–148.

35. Barboiu M & Gilles A (2013) From Natural to Bioassisted and Biomimetic Artificial Water Channel Systems. Acc. Chem. Res. 46(12):2814–2823.

36. Barboiu M (2012) Artificial water channels. Angew. Chem. Int. Ed. Engl. 51(47):11674–11676.

37. Le Duc Y, et al. (2011) Imidazole-Quartet Water and Proton Dipolar Channels. Angew. Chem. Int. Ed. 50(48):11366–11372.

38. Beitz E, Wu B, Holm LM, Schultz JE, & Zeuthen T (2006) Point mutations in the aromatic/arginine region in aquaporin 1 allow passage of urea, glycerol, ammonia, and protons. Proc Natl Acad Sci U S A 103(2):269–274.

39. de Groot BL & Grubmuller H (2005) The dynamics and energetics of water permeation and proton exclusion in aquaporins. Curr. Opin. Struct. Biol. 15(2):176–183.

40. Li H, et al. (2011) Enhancement of Proton Conductance by Mutations of the Selectivity Filter of Aquaporin-1. J. Mol. Biol. 407(4):607–620.

41. Lee JK, et al. (2005) Structural basis for conductance by the archaeal aquaporin AqpM at 1.68 A. Proc Natl Acad Sci U S A 102(52):18932–18937.

42. Savage DF, Egea PF, Robles-Colmenares Y, O’Connell JD, & Stroud RM (2003) Architecture and selectivity in aquaporins:2.5 angstrom X-ray structure of aquaporin Z. PLoS Biol. 1(3):334–340.

43. Fu D, et al. (2000) Structure of a glycerol-conducting channel and the basis for its selectivity. Science 290(5491):481–486.

44. Eriksson UK, et al. (2013) Subangstrom resolution X-ray structure details aquaporin-water interactions. Science 340(6138):1346–1349.

45. Newby ZE, et al. (2008) Crystal structure of the aquaglyceroporin PfAQP from the malarial parasite Plasmodium falciparum. Nat. Struct. Mol. Biol. 15(6):619–625.

46. Gonen T, et al. (2005) Lipid-protein interactions in double-layered two-dimensional AQP0 crystals. Nature 438(7068):633–638.

47. Harries WE, Akhavan D, Miercke LJ, Khademi S, & Stroud RM (2004) The channel architecture of aquaporin 0 at a 2.2-A resolution. Proc Natl Acad Sci U S A 101(39):14045–14050.

48. Carrillo DR, et al. (2014) Crystallization and preliminary crystallographic analysis of human aquaporin 1 at a resolution of 3.28 angstrom. Acta Crystallographica Section F-Structural Biology Communications 70:1657–1663.

49. Sui HX, Han BG, Lee JK, Walian P, & Jap BK (2001) Structural basis of water-specific transport through the AQP1 water channel. Nature 414(6866):872–878.

50. Frick A, et al. (2014) X-ray structure of human aquaporin 2 and its implications for nephrogenic diabetes insipidus and trafficking. Proc Natl Acad Sci U S A 111(17):6305–6310.

51. Ho JD, et al. (2009) Crystal structure of human aquaporin 4 at 1.8 A and its mechanism of conductance. Proc Natl Acad Sci U S A 106(18):7437–7442.

52. Tani K, et al. (2009) Mechanism of aquaporin-4’s fast and highly selective water conduction and proton exclusion. J. Mol. Biol. 389(4):694–706.

53. Horsefield R, et al. (2008) High-resolution x-ray structure of human aquaporin 5. Proc Natl Acad Sci U S A 105(36):13327–13332.

54. de Mare SW, Venskutonyte R, Eltschkner S, de Groot BL, & Lindkvist-Petersson K (2020) Structural Basis for Glycerol Efflux and Selectivity of Human Aquaporin 7. Structure 28(2):215–222 e213.

55. Gotfryd K, et al. (2018) Human adipose glycerol flux is regulated by a pH gate in AQP10. Nat Commun 9(1):4749.

56. Tornroth-Horsefield S, et al. (2006) Structural mechanism of plant aquaporin gating. Nature 439(7077):688–694.

57. Wang H, Schoebel S, Schmitz F, Dong H, & Hedfalk K (2020) Characterization of aquaporin-driven hydrogen peroxide transport. Biochim Biophys Acta Biomembr 1862(2):183065.

58. Kirscht A, et al. (2016) Crystal Structure of an Ammonia-Permeable Aquaporin. PLoS Biol. 14(3):e1002411.

59. Bienert GP, et al. (2007) Specific aquaporins facilitate the diffusion of hydrogen peroxide across membranes. J. Biol. Chem. 282(2):1183–1192.

60. Hub JS & de Groot BL (2008) Mechanism of selectivity in aquaporins and aquaglyceroporins. Proc Natl Acad Sci U S A 105(4):1198–1203.

61. Wang Y, Schulten K, & Tajkhorshid E (2005) What makes an aquaporin a glycerol channel? A comparative study of AqpZ and GlpF. Structure 13(8):1107–1118.

62. Savage DF, O’Connell JD, 3rd, Miercke LJ, Finer-Moore J, & Stroud RM (2010) Structural context shapes the aquaporin selectivity filter. Proc Natl Acad Sci U S A 107(40):17164–17169.

63. Oliva R, Calamita G, Thornton JM, & Pellegrini-Calace M (2010) Electrostatics of aquaporin and aquaglyceroporin channels correlates with their transport selectivity. Proc Natl Acad Sci U S A 107(9):4135–4140.

64. Kitchen P, et al. (2019) Water channel pore size determines exclusion properties but not solute selectivity. Scientific Reports 9(1):20369.

65. Wu B & Beitz E (2007) Aquaporins with selectivity for unconventional permeants. Cell. Mol. Life Sci. 64(18):2413–2421.

66. Bienert GP & Chaumont F (2014) Aquaporin-facilitated transmembrane diffusion of hydrogen peroxide. Biochim. Biophys. Acta 1840(5):1596–1604.

67. Mukhopadhyay R & Beitz E (2010) Metalloid transport by aquaglyceroporins: consequences in the treatment of human diseases. Adv. Exp. Med. Biol. 679:57–69.

68. Bienert GP, Schussler MD, & Jahn TP (2008) Metalloids: essential, beneficial or toxic? Major intrinsic proteins sort it out. Trends Biochem. Sci. 33(1):20–26.

69. Rothert M, Ronfeldt D, & Beitz E (2017) Electrostatic attraction of weak monoacid anions increases probability for protonation and passage through aquaporins. J. Biol. Chem. 292(22):9358–9364.

70. Schmidt JDR, Walloch P, Hoger B, & Beitz E (2021) Aquaporins with lactate/lactic acid permeability at physiological pH conditions. Biochimie.

71. Bienert GP, Desguin B, Chaumont F, & Hols P (2013) Channel-mediated lactic acid transport: a novel function for aquaglyceroporins in bacteria. Biochem. J. 454(3):559–570.

72. Mori IC, et al. (2014) CO2 Transport by PIP2 Aquaporins of Barley. Plant and Cell Physiology 55(2):251–257.

73. Montiel V, et al. (2020) Inhibition of aquaporin-1 prevents myocardial remodeling by blocking the transmembrane transport of hydrogen peroxide. Science Translational Medicine 12(564):eaay2176.

74. Boytsov D, Hannesschlaeger C, Horner A, Siligan C, & Pohl P (2020) Micropipette Aspiration-Based Assessment of Single Channel Water Permeability. Biotechnol J 15(7):e1900450.

75. Hannesschlaeger C, Horner A, & Pohl P (2019) Intrinsic Membrane Permeability to Small Molecules. Chem. Rev. 119(9):5922–5953.

76. Hannesschlager C, Barta T, Siligan C, & Horner A (2018) Quantification of Water Flux in Vesicular Systems. Sci Rep 8(1):8516.

77. Horner A & Pohl P (2018) Single-file transport of water through membrane channels. Faraday Discuss. 209(0):9–33.

78. Wachlmayr J, et al. (2021) Scattering versus Fluorescence Self-Quenching: More than a Question of Faith for the Quantification of Water Flux in Large Unilamellar Vesicles? Nanoscale Advances.

79. Hoomann T, Jahnke N, Horner A, Keller S, & Pohl P (2013) Filter gate closure inhibits ion but not water transport through potassium channels. Proc Natl Acad Sci U S A 110(26):10842–10847.

80. Knyazev DG, et al. (2013) The bacterial translocon SecYEG opens upon ribosome binding. J. Biol. Chem. 288(25):17941–17946.

81. Horner A, Siligan C, Cornean A, & Pohl P (2018) Positively charged residues at the channel mouth boost single-file water flow. Faraday Discuss. 209(0):55–65.

82. Gravelle S, et al. (2013) Optimizing water permeability through the hourglass shape of aquaporins. Proc Natl Acad Sci U S A 110(41):16367–16372.

83. Horner A & Pohl P (2018) Comment on “Enhanced water permeability and tunable ion selectivity in subnanometer carbon nanotube porins”. Science 359(6383).

84. Erokhova L, Horner A, Kugler P, & Pohl P (2011) Monitoring single-channel water permeability in polarized cells. J. Biol. Chem. 286(46):39926–39932.

85. Zardoya R (2005) Phylogeny and evolution of the major intrinsic protein family. Biol. Cell 97(6):397–414.

86. Finn RN & Cerda J (2015) Evolution and functional diversity of aquaporins. Biol. Bull. 229(1):6–23.

87. Hedfalk K, et al. (2006) Aquaporin gating. Curr. Opin. Struct. Biol. 16(4):447–456.

88. Tornroth-Horsefield S, Hedfalk K, Fischer G, Lindkvist-Petersson K, & Neutze R (2010) Structural insights into eukaryotic aquaporin regulation. FEBS Lett. 584(12):2580–2588.

89. Frick A, Jarva M, & Tornroth-Horsefield S (2013) Structural basis for pH gating of plant aquaporins. FEBS Lett. 587(7):989–993.

90. Sachdeva R & Singh B (2014) Insights into structural mechanisms of gating induced regulation of aquaporins. Prog. Biophys. Mol. Biol. 114(2):69–79.

91. Bezerra-Neto JP, et al. (2019) Plant Aquaporins: Diversity, Evolution and Biotechnological Applications. Curr Protein Pept Sci 20(4):368–395.

92. del Carmen Martinez-Ballesta M & Carvajal M (2014) New challenges in plant aquaporin biotechnology. Plant Sci. 217:71–77.

93. Gena P, Pellegrini-Calace M, Biasco A, Svelto M, & Calamita G (2011) Aquaporin Membrane Channels: Biophysics, Classification, Functions, and Possible Biotechnological Applications. Food Biophysics 6(2):241–249.

94. Roche JV & Tornroth-Horsefield S (2017) Aquaporin Protein-Protein Interactions. Int J Mol Sci 18(11):2255.

95. Sjöhamn J & Hedfalk K (2014) Unraveling aquaporin interaction partners. Biochimica et Biophysica Acta (BBA)-General Subjects 1840(5):1614–1623.

96. Strugatsky D, et al. (2013) Structure of the proton-gated urea channel from the gastric pathogen Helicobacter pylori. Nature 493(7431):255–258.

97. Cui Y, et al. (2019) pH-dependent gating mechanism of the Helicobacter pylori urea channel revealed by cryo-EM. Sci Adv 5(3):eaav8423.

98. Fetter K, Van Wilder V, Moshelion M, & Chaumont F (2004) Interactions between plasma membrane aquaporins modulate their water channel activity. Plant Cell 16(1):215–228.

99. Berny MC, Gilis D, Rooman M, & Chaumont F (2016) Single mutations in the transmembrane domains of maize plasma membrane aquaporins affect the activity of monomers within a heterotetramer. Molecular plant 9(7):986–1003.

100. Heinen RB, et al. (2014) Expression and characterization of plasma membrane aquaporins in stomatal complexes of Zea mays. Plant Mol. Biol. 86(3):335–350.

101. Jozefkowicz C, et al. (2016) PIP Water Transport and Its pH Dependence Are Regulated by Tetramer Stoichiometry. Biophys. J. 110(6):1312–1321.

102. Hiroaki Y, et al. (2006) Implications of the aquaporin-4 structure on array formation and cell adhesion. J. Mol. Biol. 355(4):628–639.

103. Yang B, Brown D, & Verkman AS (1996) The mercurial insensitive water channel (AQP-4) forms orthogonal arrays in stably transfected Chinese hamster ovary cells. J. Biol. Chem. 271(9):4577–4580.

104. Rossi A, Moritz TJ, Ratelade J, & Verkman AS (2012) Super-resolution imaging of aquaporin-4 orthogonal arrays of particles in cell membranes. J. Cell Sci. 125(Pt 18):4405–4412.

105. Silberstein C, et al. (2004) Membrane organization and function of M1 and M23 isoforms of aquaporin-4 in epithelial cells. Am J Physiol Renal Physiol 287(3):F501–511.

106. Fenton RA, et al. (2010) Differential water permeability and regulation of three aquaporin 4 isoforms. Cell. Mol. Life Sci. 67(5):829–840.

107. Zhang H & Verkman AS (2008) Evidence against involvement of aquaporin-4 in cell-cell adhesion. J. Mol. Biol. 382(5):1136–1143.

108. Frydenlund DS, et al. (2006) Temporary loss of perivascular aquaporin-4 in neocortex after transient middle cerebral artery occlusion in mice. Proc Natl Acad Sci U S A 103(36):13532–13536.

109. Noell S, Fallier-Becker P, Deutsch U, Mack AF, & Wolburg H (2009) Agrin defines polarized distribution of orthogonal arrays of particles in astrocytes. Cell Tissue Res. 337(2):185–195.

110. Saparov SM, Kozono D, Rothe U, Agre P, & Pohl P (2001) Water and ion permeation of aquaporin-1 in planar lipid bilayers. Major differences in structural determinants and stoichiometry. J. Biol. Chem. 276(34):31515–31520.

111. Tsunoda SP, Wiesner B, Lorenz D, Rosenthal W, & Pohl P (2004) Aquaporin-1, nothing but a water channel. J. Biol. Chem. 279(12):11364–11367.

112. Agmon N (1995) The Grotthuss Mechanism. Chem. Phys. Lett. 244(5-6):456–462.

113. Ballesteros JA & Weinstein H (1995) [19] Integrated methods for the construction of three-dimensional models and computational probing of structure-function relations in G protein-coupled receptors. Receptor Molecular Biology, Methods in Neurosciences, (Elsevier), Vol 25, pp 366–428.

114. von Heijne G (1992) Membrane protein structure prediction. Hydrophobicity analysis and the positive-inside rule. J. Mol. Biol. 225(2):487–494.

115. Fischer G, et al. (2009) Crystal structure of a yeast aquaporin at 1.15 angstrom reveals a novel gating mechanism. PLoS Biol. 7(6):e1000130.

116. Kozono D, et al. (2003) Functional expression and characterization of an archaeal aquaporin. AqpM from methanothermobacter marburgensis. J. Biol. Chem. 278(12):10649–10656.

117. Ishibashi K, Morishita Y, & Tanaka Y (2017) The Evolutionary Aspects of Aquaporin Family. Aquaporins, ed Yang B (Springer Netherlands, Dordrecht), pp 35–50.

118. Zardoya R, Ding X, Kitagawa Y, & Chrispeels MJ (2002) Origin of plant glycerol transporters by horizontal gene transfer and functional recruitment. Proc Natl Acad Sci U S A 99(23):14893–14896.

119. Klenk HP, et al. (1997) The complete genome sequence of the hyperthermophilic, sulphate-reducing archaeon Archaeoglobus fulgidus. Nature 390(6658):364–370.

120. Lamosa P, et al. (2000) Thermostabilization of proteins by diglycerol phosphate, a new compatible solute from the hyperthermophile Archaeoglobus fulgidus. Appl. Environ. Microbiol. 66(5):1974–1979.

121. Martins LO, et al. (1997) Organic solutes in hyperthermophilic archaea. Appl. Environ. Microbiol. 63(3):896–902.

122. Smart OS, Goodfellow JM, & Wallace BA (1993) The pore dimensions of gramicidin A. Biophys. J. 65(6):2455–2460.

123. Smart OS, Neduvelil JG, Wang X, Wallace BA, & Sansom MS (1996) HOLE: a program for the analysis of the pore dimensions of ion channel structural models. J Mol Graph 14(6):354–360, 376.

124. Böhm H-J, Brode S, Hesse U, & Klebe G (1996) Oxygen and Nitrogen in Competitive Situations: Which is the Hydrogen-Bond Acceptor? Chemistry - A European Journal 2(12):1509–1513.

125. Sterpone F, Stirnemann G, Hynes JT, & Laage D (2010) Water hydrogen-bond dynamics around amino acids: the key role of hydrophilic hydrogen-bond acceptor groups. J. Phys. Chem. B 114(5):2083–2089.

126. Smolin N, Li B, Beck DA, & Daggett V (2008) Side-chain dynamics are critical for water permeation through aquaporin-1. Biophys. J. 95(3):1089–1098.

127. McNulty R, Ulmschneider JP, Luecke H, & Ulmschneider MB (2013) Mechanisms of molecular transport through the urea channel of Helicobacter pylori. Nat Commun 4:2900.

128. Verdoucq L, Grondin A, & Maurel C (2008) Structure–function analysis of plant aquaporin At PIP2; 1 gating by divalent cations and protons. Biochem. J. 415(3):409–416.

129. Rodrigues C, et al. (2016) Rat Aquaporin-5 Is pH-Gated Induced by Phosphorylation and Is Implicated in Oxidative Stress. Int J Mol Sci 17(12).

130. Reichow SL, et al. (2013) Allosteric mechanism of water-channel gating by Ca2+–calmodulin. Nat. Struct. Mol. Biol. 20:1085.

131. Xin L, et al. (2011) Water permeation dynamics of AqpZ: a tale of two states. Biochim. Biophys. Acta 1808(6):1581–1586.

132. Zhu F, Tajkhorshid E, & Schulten K (2001) Molecular dynamics study of aquaporin-1 water channel in a lipid bilayer. FEBS Lett. 504(3):212–218.

133. Mom R, et al. (2021) Voltage-gating of aquaporins, a putative conserved safety mechanism during ionic stresses. FEBS Lett. 595(1):41–57.

134. Hub JS, Aponte-Santamaria C, Grubmuller H, & de Groot BL (2010) Voltage-Regulated Water Flux through Aquaporin Channels In Silico. Biophys. J. 99(12):L97–L99.

135. Murata K, et al. (2000) Structural determinants of water permeation through aquaporin-1. Nature 407(6804):599–605.

136. Jiang J, Daniels BV, & Fu D (2006) Crystal structure of AqpZ tetramer reveals two distinct Arg-189 conformations associated with water permeation through the narrowest constriction of the water-conducting channel. J. Biol. Chem. 281(1):454–460.

137. Zhao Y, et al. (2018) Gating Mechanism of Aquaporin Z in Synthetic Bilayers and Native Membranes Revealed by Solid-State NMR Spectroscopy. J. Am. Chem. Soc. 140(25):7885–7895.

138. Heiby JC, Goretzki B, Johnson CM, Hellmich UA, & Neuweiler H (2019) Methionine in a protein hydrophobic core drives tight interactions required for assembly of spider silk. Nature Communications 10(1):4378.

139. Janosi L & Ceccarelli M (2013) The gating mechanism of the human aquaporin 5 revealed by molecular dynamics simulations. PLoS One 8(4):e59897.

140. Alberga D, et al. (2014) A new gating site in human aquaporin-4: Insights from molecular dynamics simulations. Biochim. Biophys. Acta 1838(12):3052–3060.

141. Kaptan S, et al. (2015) H95 Is a pH-Dependent Gate in Aquaporin 4. Structure 23(12):2309–2318.

142. Nemeth-Cahalan KL & Hall JE (2000) pH and calcium regulate the water permeability of aquaporin 0. J. Biol. Chem. 275(10):6777–6782.

143. Nemeth-Cahalan KL, Kalman K, & Hall JE (2004) Molecular basis of pH and Ca2+ regulation of aquaporin water permeability. J. Gen. Physiol. 123(5):573–580.

144. Soto G, et al. (2010) TIP5;1 is an aquaporin specifically targeted to pollen mitochondria and is probably involved in nitrogen remobilization in Arabidopsis thaliana. Plant J. 64(6):1038–1047.

145. Zeuthen T & Klaerke DA (1999) Transport of water and glycerol in aquaporin 3 is gated by H+. J. Biol. Chem. 274(31):21631–21636.

146. Zelenina M, Bondar AA, Zelenin S, & Aperia A (2003) Nickel and extracellular acidification inhibit the water permeability of human aquaporin-3 in lung epithelial cells. J. Biol. Chem. 278(32):30037–30043.

147. de Almeida A, et al. (2016) Exploring the gating mechanisms of aquaporin-3: new clues for the design of inhibitors? Mol Biosyst 12(5):1564–1573.

148. Tournaire-Roux C, et al. (2003) Cytosolic pH regulates root water transport during anoxic stress through gating of aquaporins. Nature 425(6956):393–397.

149. Yasui M, et al. (1999) Rapid gating and anion permeability of an intracellular aquaporin. Nature 402(6758):184–187.

150. Mosca AF, et al. (2018) Molecular Basis of Aquaporin-7 Permeability Regulation by pH. Cells 7(11):207.

151. Leitão L, Prista C, Moura TF, Loureiro-Dias MC, & Soveral G (2012) Grapevine aquaporins: gating of a tonoplast intrinsic protein (TIP2; 1) by cytosolic pH. PLoS One 7(3):e33219.

152. Fischer M & Kaldenhoff R (2008) On the pH regulation of plant aquaporins. J. Biol. Chem. 283(49):33889–33892.

153. Olsson MH, Sondergaard CR, Rostkowski M, & Jensen JH (2011) PROPKA3: Consistent Treatment of Internal and Surface Residues in Empirical pKa Predictions. J Chem Theory Comput 7(2):525–537.

154. Søndergaard CR, Olsson MH, Rostkowski M, & Jensen JH (2011) Improved treatment of ligands and coupling effects in empirical calculation and rationalization of p K a values. Journal of chemical theory and computation 7(7):2284–2295.

155. Truelsen SF, et al. (2021) The role of water coordination in the pH-dependent gating of hAQP10. Biochim Biophys Acta Biomembr 1864(1):183809.

156. Steinbrecher T, et al. (2012) Peptide-lipid interactions of the stress-response peptide TisB that induces bacterial persistence. Biophys. J. 103(7):1460–1469.

157. Bahadur RP, Chakrabarti P, Rodier F, & Janin J (2003) Dissecting subunit interfaces in homodimeric proteins. Proteins 53(3):708–719.

158. Xu D, Tsai CJ, & Nussinov R (1997) Hydrogen bonds and salt bridges across protein-protein interfaces. Protein Eng. 10(9):999–1012.

159. Gupta K, et al. (2017) The role of interfacial lipids in stabilizing membrane protein oligomers. Nature 541(7637):421–424.

160. Krissinel E & Henrick K (2007) Inference of macromolecular assemblies from crystalline state. J. Mol. Biol. 372(3):774–797.

161. Lomize MA, Lomize AL, Pogozheva ID, & Mosberg HI (2006) OPM: orientations of proteins in membranes database. Bioinformatics 22(5):623–625.

162. McDonald IK & Thornton JM (1994) Satisfying hydrogen bonding potential in proteins. J. Mol. Biol. 238(5):777–793.

163. Pace CN, Shirley BA, McNutt M, & Gajiwala K (1996) Forces contributing to the conformational stability of proteins. FASEB J. 10(1):75–83.

164. Fersht AR (1987) The Hydrogen-Bond in Molecular Recognition. Trends Biochem. Sci. 12(8):301–304.

165. Horovitz A, Serrano L, Avron B, Bycroft M, & Fersht AR (1990) Strength and co-operativity of contributions of surface salt bridges to protein stability. J. Mol. Biol. 216(4):1031–1044.

166. Akke M & Forsen S (1990) Protein Stability and Electrostatic Interactions between Solvent Exposed Charged Side-Chains. Proteins-Structure Function and Genetics 8(1):23–29.

167. Betz SF (1993) Disulfide bonds and the stability of globular proteins. Protein Sci. 2(10):1551–1558.

168. Clarke J & Fersht AR (1993) Engineered disulfide bonds as probes of the folding pathway of barnase: increasing the stability of proteins against the rate of denaturation. Biochemistry 32(16):4322–4329.

169. Kim S, et al. (2005) Transmembrane glycine zippers: physiological and pathological roles in membrane proteins. Proc Natl Acad Sci U S A 102(40):14278–14283.

170. Situ AJ, et al. (2018) Membrane Anchoring of alpha-Helical Proteins: Role of Tryptophan. J. Phys. Chem. B 122(3):1185–1194.

171. Laganowsky A, et al. (2014) Membrane proteins bind lipids selectively to modulate their structure and function. Nature 510(7503):172–175.

172. Schmidt V, Sidore M, Bechara C, Duneau JP, & Sturgis JN (2019) The lipid environment of Escherichia coli Aquaporin Z. Biochim Biophys Acta Biomembr 1861(2):431–440.

173. Corey RA, et al. (2021) Identification and assessment of cardiolipin interactions with E. coli inner membrane proteins. Sci Adv 7(34):eabh2217.

174. Pluhackova K & Horner A (2021) Native-like membrane models of E. coli polar lipid extract shed light on the importance of lipid composition complexity. BMC Biol. 19(1):4.

175. Garg SG & Gould SB (2016) The Role of Charge in Protein Targeting Evolution. Trends Cell Biol. 26(12):894–905.

176. von Heijne G (1989) Control of topology and mode of assembly of a polytopic membrane protein by positively charged residues. Nature 341(6241):456–458.

177. Neumann J, Klein N, Otzen DE, & Schneider D (2014) Folding energetics and oligomerization of polytopic alpha-helical transmembrane proteins. Arch. Biochem. Biophys. 564:281–296.

178. Yool AJ & Weinstein AM (2002) New roles for old holes: Ion channel function in aquaporin-1. News in Physiological Sciences 17:68–72.

179. Yool AJ & Campbell EM (2012) Structure, function and translational relevance of aquaporin dual water and ion channels. Molecular Aspects of Medicine 33(5):553–561.

180. Tyerman SD, McGaughey SA, Qiu J, Yool AJ, & Byrt CS (2021) Adaptable and Multifunctional Ion-Conducting Aquaporins. Annu. Rev. Plant Biol. 72(1):703–736.

181. Kourghi M, et al. (2018) Fundamental structural and functional properties of Aquaporin ion channels found across the kingdoms of life. Clin. Exp. Pharmacol. Physiol. 45(4):401–409.

182. Wang Y, Cohen J, Boron WF, Schulten K, & Tajkhorshid E (2007) Exploring gas permeability of cellular membranes and membrane channels with molecular dynamics. J Struct Biol 157(3):534–544.

183. Portella G & de Groot BL (2009) Determinants of water permeability through nanoscopic hydrophilic channels. Biophys. J. 96(3):925–938.

184. Berezhkovskii A & Hummer G (2002) Single-file transport of water molecules through a carbon nanotube. Phys. Rev. Lett. 89(6):064503.

185. Stroud RM, Nollert P, & Miercke L (2003) The glycerol facilitator GlpF, its aquaporin family of channels, and their selectivity. Membrane Proteins 63:291–316.

186. Cymer F & Schneider D (2010) A single glutamate residue controls the oligomerization, function, and stability of the aquaglyceroporin GlpF. Biochemistry 49(2):279–286.

187. Borgnia MJ & Agre P (2001) Reconstitution and functional comparison of purified GlpF and AqpZ, the glycerol and water channels from Escherichia coli. Proc Natl Acad Sci U S A 98(5):2888–2893.

188. Galka JJ, Baturin SJ, Manley DM, Kehler AJ, & O’neil JD (2008) Stability of the glycerol facilitator in detergent solutions. Biochemistry 47(11):3513–3524.

189. Doyle DA, et al. (1998) The structure of the potassium channel: molecular basis of K+ conduction and selectivity. Science 280(5360):69–77.

190. Payandeh J, Scheuer T, Zheng N, & Catterall WA (2011) The crystal structure of a voltage-gated sodium channel. Nature 475(7356):353–358.

191. Neumann E & Rosenheck K (1972) Permeability changes induced by electric impulses in vesicular membranes. The Journal of Membrane Biology 10(1):279–290.

192. Kirsch SA & Bockmann RA (2016) Membrane pore formation in atomistic and coarse-grained simulations. Biochim. Biophys. Acta 1858(10):2266–2277.

193. Huang F, Fang Z, Mast J, & Chen W (2013) Comparison of membrane electroporation and protein denature in response to pulsed electric field with different durations. Bioelectromagnetics 34(4):253–263.

194. Patel R, Smith SM, & Robinson C (2014) Protein transport by the bacterial Tat pathway. Biochimica et Biophysica Acta (BIOCHIMICA ET BIOPHYSICA ACTA) - Molecular Cell Research 1843(8):1620–1628.

195. Klein N, Hellmann N, & Schneider D (2015) Anionic Lipids Modulate the Activity of the Aquaglyceroporin GlpF. Biophys. J. 109(4):722–731.

196. Schmidt V & Sturgis JN (2017) Making Monomeric Aquaporin Z by Disrupting the Hydrophobic Tetramer Interface. ACS Omega 2(6):3017–3027.

197. Cymer F & Schneider D (2012) Oligomerization of polytopic alpha-helical membrane proteins: causes and consequences. Biol. Chem. 393(11):1215–1230.

198. Trefz M, Keller R, Vogt M, & Schneider D (2017) The GlpF residue Trp219 is part of an amino-acid cluster crucial for aquaglyceroporin oligomerization and function. Biochim. Biophys. Acta.

199. Klein N, Trefz M, & Schneider D (2019) Covalently Linking Oligomerization-Impaired GlpF Protomers Does Not Completely Re-establish Wild-Type Channel Activity. International Journal of Molecular Sciences 20(4):927.

200. Sholl DS & Lively RP (2016) Seven chemical separations: to change the world: purifying mixtures without using heat would lower global energy use, emissions and pollution--and open up new routes to resources. Nature 532(7600):435–438.

201. Kumar M, Grzelakowski M, Zilles J, Clark M, & Meier W (2007) Highly permeable polymeric membranes based on the incorporation of the functional water channel protein Aquaporin Z. Proc Natl Acad Sci U S A 104(52):20719–20724.

202. AQUAPORIN (2020) Aquaporin Inside.

203. Waterhouse AM, Procter JB, Martin DM, Clamp M, & Barton GJ (2009) Jalview Version 2--a multiple sequence alignment editor and analysis workbench. Bioinformatics 25(9):1189–1191.

204. Kumar M, Habel JE, Shen YX, Meier WP, & Walz T (2012) High-density reconstitution of functional water channels into vesicular and planar block copolymer membranes. J. Am. Chem. Soc. 134(45):18631–18637.

205. Krissinel E (2010) Crystal Contacts as Nature’s Docking Solutions. J. Comput. Chem. 31(1):133–143.

206. Schrödinger LLC (2020) The PyMOL Molecular Graphics System, Version~2.4.

207. R. Core Team (2020) R: A Language and Environment for Statistical Computing (Vienna, Austria, http://www.R-project.org).

208. Tamura K, Stecher G, & Kumar S (2021) MEGA11: Molecular Evolutionary Genetics Analysis Version 11. Mol. Biol. Evol. 38(7):3022–3027.

209. Abraham MJ, et al. (2015) GROMACS: High performance molecular simulations through multi-level parallelism from laptops to supercomputers. SoftwareX 1-2:19–25.

210. Pluhackova KH, Andreas (2020) CHARMM36 and Martini3 membrane models of E. coli polar lipid extract shed light on the importance of lipid composition complexity. BMC Biology.

211. Pluhackova K, Wassenaar TA, & Bockmann RA (2013) Molecular dynamics simulations of membrane proteins. Methods Mol Biol 1033:85–101.

212. Hu G, Qi L, Dou X, & Wang J (2013) The influences of protonation state of histidine on aromatic/arginine region of aquaporin-1 protein. Molecular Simulation 39(4):261–269.

213. Bjelkmar P, Larsson P, Cuendet MA, Hess B, & Lindahl E (2010) Implementation of the CHARMM Force Field in GROMACS: Analysis of Protein Stability Effects from Correction Maps, Virtual Interaction Sites, and Water Models. Journal of chemical theory and computation 6(2):459–466.

214. Huang J, et al. (2017) CHARMM36m: an improved force field for folded and intrinsically disordered proteins. Nature Methods 14(1):71–73.

215. Klauda JB, et al. (2010) Update of the CHARMM All-Atom Additive Force Field for Lipids: Validation on Six Lipid Types. The Journal of Physical Chemistry B 114(23):7830–7843.

216. Jorgensen WL & Madura JD (1985) Temperature and size dependence for Monte Carlo simulations of TIP4P water. Molecular Physics 56(6):1381–1392.

217. Beckstein O & Sansom MSP (2004) The influence of geometry, surface character, and flexibility on the permeation of ions and water through biological pores. Physical Biology 1(1):42–52.

